# Sexual antagonism evolves when autosomes influence offspring sex ratio

**DOI:** 10.1101/2023.06.14.544982

**Authors:** Solomon Sloat, Matthew Rockman

**Affiliations:** Department of Biology and Center for Genomics and Systems Biology, New York University, New York, NY 10003

## Abstract

Sex allocation theory generally assumes maternal control of offspring sex and makes few predictions for populations evolving under paternal control. Using population genetic simulations, we show that maternal and paternal control of the sex ratio lead to different equilibrium sex ratios in structured populations. Sex ratios evolved under paternal control are more female biased. This effect is dependent on the population subdivision; fewer founding individuals leads to both more biased sex ratios and a greater difference between the paternal and maternal equilibria. In addition, sexual antagonism evolves in simulations with both maternally- and paternally-acting loci. Maternally-acting loci continuously accumulate ever more female-biasing effects as male-biasing effects accumulate at paternally-acting loci. The difference in evolved sex-ratio equilibria and the evolution of sexual antagonism can be largely explained by differences in the between-group variance of maternal and paternal effects in the founding generation. These theoretical results apply to any system with biparental autosomal influence over offspring sex, opening up an exciting new line of questioning.

## Introduction

Ecological pressures on reproduction and survival shape life histories (Stearns, 1992). One fundamental life-history trait is the allocation of resources towards offspring of different sexes. Sex-ratio theory has successfully connected important ecological and genetic factors to sex-ratio evolution (Charnov, 1982; West, 2009). However, evolutionary research has neglected important aspects of sex-ratio regulation. In typical diploid species, autosomal genes that affect sex ratio can act in mothers or in fathers or in both. Yet the implications of paternally-controlled sex allocation, the possibility of differing equilibria under paternal and maternal systems, and the potential for sexual antagonism over offspring sex has never been modeled for autosomal genetic systems and has received limited experimental attention (Douhard & Geffroy, 2021; Shuker et al., 2009).

Under panmixia, frequency-dependent selection drives sex ratios to equality (Düsing, 1884; Edwards, 2000; Fisher, 1930). Mutations that produce more of the rarer sex are favored until the population reaches the unbeatable 50:50 strategy. In some classes of structured populations, however, sex ratios are predicted to evolve a female bias. Many classic models, a subset of which we discuss here, have investigated the causes and consequences of such a bias using different population structure conceptions and modeling strategies. Local Mate Competition (LMC) assumes a population subdivided into ephemeral patches where a number of founding mothers produce equal numbers of progeny, and those progeny then mate locally before dispersal (Hamilton, 1967). Given these conditions, founding mothers that produce a female-biased sex ratio maximize their number of grandchildren while reducing competition between male progeny for mates. Later models explored the possibility of multiple generations on a patch before dispersal (Bulmer & Taylor, 1980a, 1980b; Wilson & Colwell, 1981). In these models, animals either mate locally before dispersal, as in the case of LMC, or mate after dispersal in a panmictic pool. These models differ in their approach to the original LMC formulation, describing selection acting at the level of the patch rather than at the level of the individual. Patches with more female-biased sex ratios gain a population growth advantage making up a larger portion of the migrant generation (Colwell, 1981; Wilson & Colwell, 1981).

These families of sex-ratio models have been empirically tested most extensively in haplodiploid species where maternal sex allocation is generally assumed and mechanisms of paternal sex allocation remain an open question (Douhard & Geffroy, 2021; Macke et al., 2014). One compelling piece of evidence for paternal control of offspring sex comes from a study of haplodiploid spider mites (*Tetranichus urticae*) evolved under varying levels of population subdivision (Macke et al., 2014). This study supported theoretical predictions that males in arrhenotokous species (whose males develop from unfertilized haploid eggs) should always favor female-biased sex ratios, as they only pass genes to daughters, while female-optimal sex ratios should depend on the number of founding mothers on a patch (Hawkes, 1992). Antagonism in this system is greatest under panmictic conditions, and the maternal optimum becomes more female biased as population subdivision increases. An additional study found small but significant sex-ratio variation in parasitoid haplodiploid wasp (*Nasonia vitripennis*) due to paternal genetic effects (Shuker et al., 2006).

In species with chromosomal sex determination, both males and females have overt mechanisms for adjusting offspring sex. In species where males are the heterogametic sex, fathers can influence the sex ratio through differences in sperm ability or quantity between sperm carrying sex determining chromosomes (Douhard & Geffroy, 2021; Navara, 2013; Shakes et al., 2011; Winter et al., 2017). Mothers could alter the sex ratio by selecting among sperm for fertilization, selective abortions, or through biased sex-chromosome segregation in species with ZW or Z0 sex determination (Navara, 2013). Under these systems autosomal loci could shape the intensity of sex-ratio bias exerted by each sex. These autosomal mechanisms are different from sex chromosome drive, where the biasing alleles affect their own transmission and which has been modeled extensively (Hamilton, 1967; Helleu et al., 2014). Biparental autosomal control of sex ratio is also expected in species that lack sex chromosomes, including those with polygenic sex determination.

Because autosomal genes carried by males and females are equally related to mothers and fathers, in contrast to sex-ratio alleles in haplodiploids or on sex chromosomes, modelers have largely assumed that paternal and maternal autosomal control yield identical outcomes (e.g., Bulmer & Taylor, 1980b; Wilson & Colwell, 1981). Yet simple single-locus recursion equations reveal that changes in sex ratio, population size, and allele frequencies all differ between maternal- and paternal-effect models (supplementary text), raising the possibility that the sex-ratio equilibria also differ.

A promising model system for studying paternal sex allocation is the nematode genus *Caenorhabditis*. Most species in this group, which includes the well studied genetics model *C. elegans*, are gonochorists, with separate males and females, and sex determination follows an XX/X0 system. Ancestrally, these species have female-biased sex ratios that are proximally due to competition between X-bearing and X-lacking sperm (Huang et al., 2023; LaMunyon & Ward, 1997). Because they produce these sperm, males are necessarily complicit in generating the observed sex-ratio bias. Many *Caenorhabditis* species live on ephemeral resource patches with ecologies consistent with the subdivided population structure required for the evolution of female-biased sex ratios (Félix & Braendle, 2010; Ferrari et al., 2017; Richaud et al., 2018; Sloat et al., 2022). Numerous other non-*Caenorhabditis* nematode species also exhibit sex-ratio bias, which they achieve using a variety of mechanisms (Van Goor et al., 2021). These observations raise the question of whether paternal control of sex ratio yields different evolutionary predictions than maternal control, and it sets up the possibility of sexual antagonism over sex ratio. We tested these possibilities using forward-time population-genetic simulations.

### The Model

We investigated the evolution of sex ratios in populations where sex ratio is determined by autosomal loci with quantitative effects. We considered three cases: maternally acting loci, paternally acting loci, or both maternally and paternally acting loci. Simulations were carried out in SLiM (Haller & Messer, 2019) using a non-Wright-Fisher metapopulation model. Each simulation contains 200 habitat patches (Fig. 1). A fixed number of founding individuals colonize each patch. Populations on these patches grow for a fixed number of generations. Mating is random within a patch. Females can mate at most once per generation while males may mate with multiple females, and each female has a fixed number of offspring. These differences define male and female within the model. Generations are overlapping unless noted. In the final generation of the patches, a fixed number of individuals are selected at random to constitute a migrant pool, consisting of unmated male and female individuals. All other individuals in the simulation are removed, and the migrant pool individuals then become the founders of the next cycle. Patches go extinct if all individuals in the patch are the same sex. Additional simulations explore the effects of panmictic mating in the migrant pool and mating before dispersal locally as in Bulmer & Taylor (1980b).

**Figure 1.**
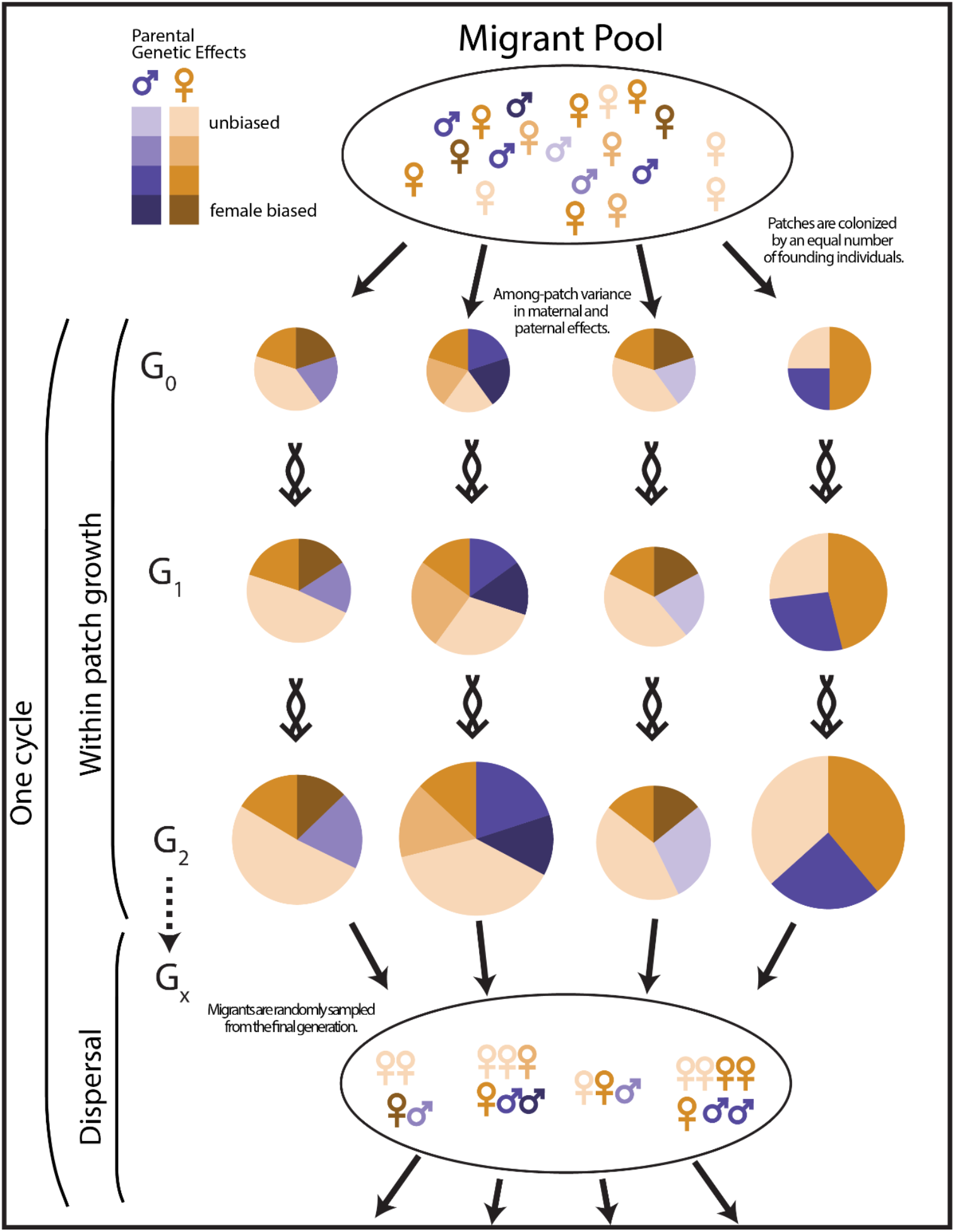
Model overview. The metapopulation is composed of many ephemeral resource patches. In the first step, founding individuals are distributed with equal numbers to each patch. Patches then grow for one or more generations as a function of a fixed brood size and the composition of the founder*’*s genotypes and sexes. Mutations arise spontaneously in offspring and may act maternally or paternally and may be male- or female-biasing. Frequency-dependent selection favors an equality of the sexes during within-patch growth, but patches with female-biased broods have a population growth advantage. In the final generation, individuals are randomly selected for the migrant pool. All other individuals are removed from the simulation and the cycle repeats itself. Patches which grow faster are overrepresented in the migrant pool. Greater between-patch variance in sex-ratio genotypes increases selection for female-biased sex ratios. Between-patch genotypic variance may be different among mothers and fathers in the founding generation which may lead to differences in selective pressures for maternal and paternal effects. All steps occur synchronously for each patch. Population growth and selection on genotypes are not drawn to scale.

Each individual has a 1 Mb long diploid genome split evenly into 5 chromosomes with a uniform recombination rate across each chromosome of 3×10^−6^. Offspring of each cross have a binomial probability of being male, and simulations start with this probability set to 0.5. Each base in the genome can mutate to affect the binomial probability, with per-base-pair mutation rate 5×10^−9^. Mutational effects are drawn with equal probability from four classes: paternal- or maternal-effect mutations that increase or decrease the binomial probability of offspring maleness. Mutations additively affect the binomial probability with a fixed effect size of 0.025. The sum of effects is unlimited within an individual but the phenotype from a mating is constrained between 1, all males, and 0, all females. Sex ratios and maternal genetic effects in females and paternal genetic effects in males are recorded every 96 generations in the first generation produced after migration. Variance of maternal and paternal effects within and across patches of the founders is also recorded every 96 generations in the founding generation. Simulations are run for 2,500-5,000 generations depending on the simulation. All scripts and data are available in the supplementary_files folder.

## Results

### Evolved sex ratios reach different equilibria for each condition

We performed simulations for each of three conditions representing three different autosomal modes of regulating sex allocation: maternal effect, paternal effect, or both simultaneously, across a range of founder numbers. As expected, female-biased sex ratios evolved in every case. Evolved sex ratio varied significantly by simulation condition (*p* < 10^−15^; analysis of deviance from a linear regression). Across all founder numbers tested, sex-ratio equilibria in simulations with paternally-acting loci were more female biased than those in simulations with maternally-acting loci (Fig. 2A,B). In simulations with both maternally- and paternally-acting loci, the evolved equilibrium was intermediate, though closer to the maternal condition (Fig. 2 A,B). The rate of evolution was different between the three conditions, with the maternal-only condition slower to evolve than the other two conditions (Fig. 2A). As expected from theory, the evolved sex ratios approach 0.5 as the number of founders in our simulations increases (Fig. 2B). We also see the differences between conditions shrink as the number of founders increases. Note that a small brood size constrains sex-ratio equilibria. This is most apparent in the two-founder paternal case where the equilibrium for a brood size of 10 is 0.228 (95% CI [0.219 0.237]) and the equilibrium for a brood size of 3 is 0.246 (95% CI [0.235 0.258]), although this is more generally true across the range of founder numbers (*p* < 10^−15^; analysis of deviance from a linear regression).

**Figure 2.**
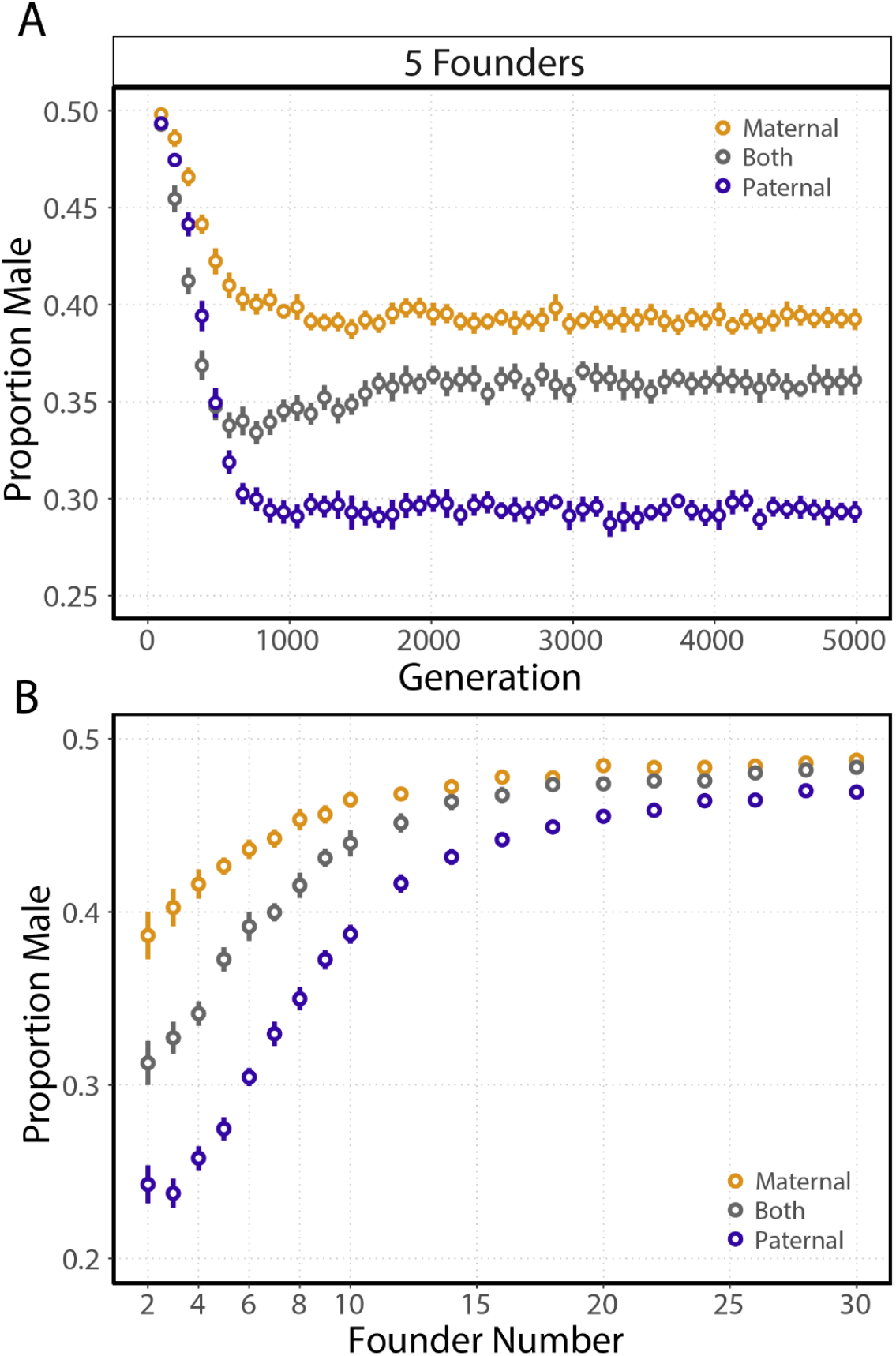
Simulations with maternally acting, paternally acting, and both maternally- and paternally-acting loci evolve different equilibrium sex ratios. **(A)** Points represent mean population sex ratios measured in the first generation after patch founding at 96-generation intervals (bars represent the 95% CI of 25 simulations; founders = 5, brood size = 10, generations before migration = 3). **(B)** Evolved sex ratios as a function of founder number. Points represent the mean population sex ratio measured in the final patch cycle, generation 4989, for the three conditions. (bars represent the 95% CI of 25 simulations; brood size = 3, generations before migration = 3).

### Sex ratio loci are sexually antagonistic

When both maternally- and paternally-acting loci are segregating in a population there is the possibility for the evolution of sexual antagonism. If the two classes of parental loci are evolving cooperatively, we would expect them to reach the same equilibrium genetic effects. Conversely, we would expect the genetic effects of the classes to evolve in opposing directions without limit if they were sexually antagonistic (Parker, 1979). We sampled maternal and paternal genetic effects for the female and male populations respectively for the first generation after founding, although males and females also carry the other sex*’*s unexpressed loci. The isolated genetic effects show clear evidence of sexual antagonism (Fig. 3A). The genetic effect of maternally-acting loci continually becomes more male biasing as male-biasing alleles are fixed over female-biasing alleles. At the same time, the genetic effect of paternally-acting loci accumulates in the other direction, favoring female bias. This stabilizes the overall population sex-ratio equilibrium. Curiously, we see that sexual antagonism evolves most rapidly at 6 founders (Fig. 3B). The genetic effects of the two classes approach the population sex ratio as the number of founders increases (Fig. 3B). In addition, antagonistic effects evolve more slowly as the number of founders increases.

**Figure 3.**
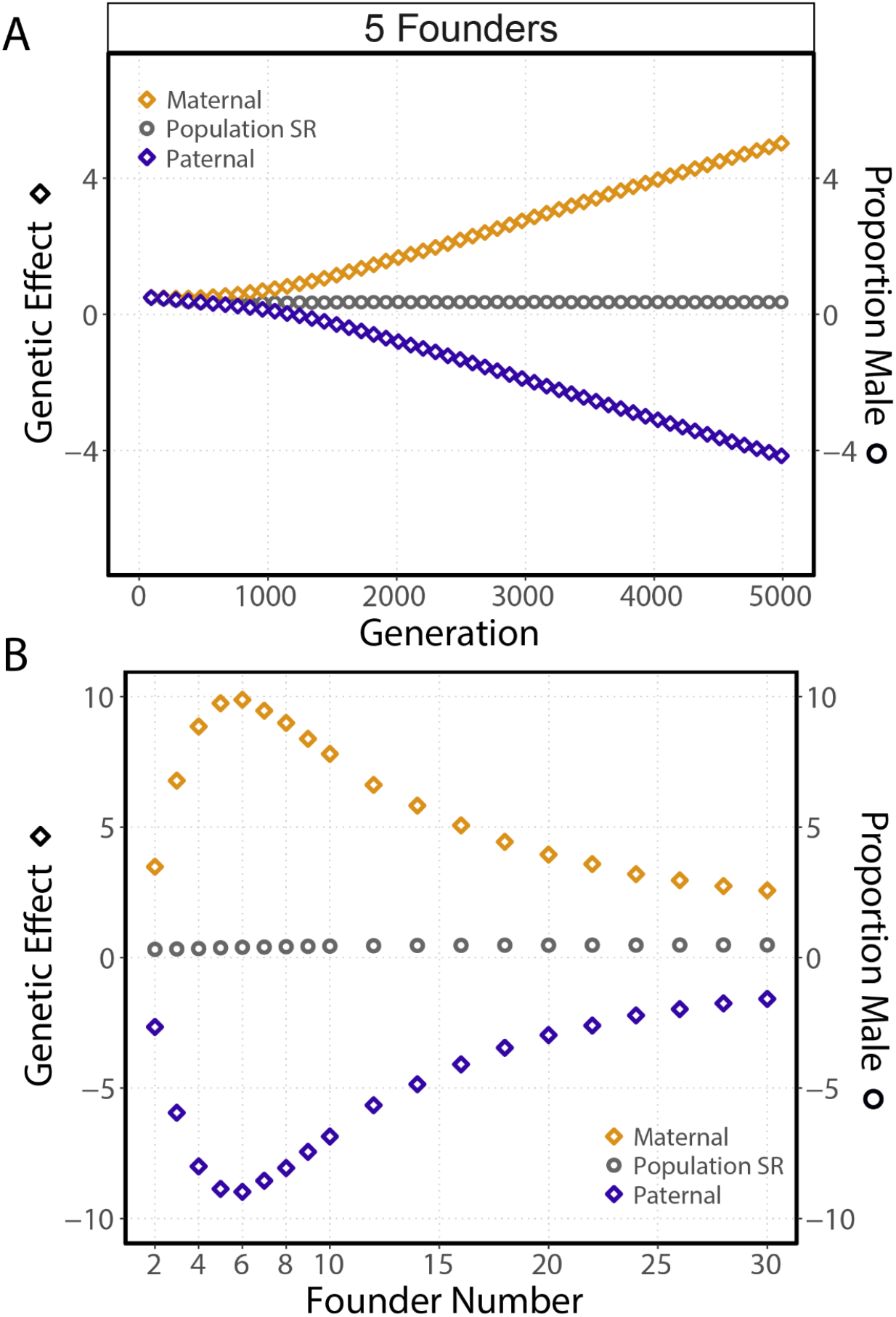
In simulations with both maternally- and paternally-acting loci, sexual antagonism evolves. **(A)** The genetic effects of maternally-acting loci and paternally-acting loci continuously evolve in opposing directions without limit. Points represent the mean genetic effects (diamonds) or population sex ratio (circles) measured in the first generation after patch founding at 96 generation intervals (bars represent the 95% CI of 25 simulations; founders = 5, brood size = 10, generations before migration = 3). **(B)** Additive genetic effect of maternally- and paternally-acting genetic effects as a function of founder number. Points represent the mean genetic effects measured in the final patch cycle, generation 4989 (bars represent the 95% CI of 25 simulations; brood size = 3, generations before migration = 3). The rate of sexual antagonism evolution (equilibrium genetic effects are never reached) is dependent on the founder number. It reaches a maximum at 6 founders and decreases as the number of founders increases.

### The effects of overlapping generations

An important metapopulation parameter is generational overlap. When generations overlap, founding individuals are able to mate with their offspring. We hypothesized that by producing more female-biased broods, founding fathers increase the probability of their own successful mating in subsequent generations. To test this hypothesis, we performed simulations over a range of founder numbers with overlapping and discrete generations. We found that the effects of overlapping generations on the equilibrium sex ratios in simulations with only maternal- or paternal-acting loci and the evolution of sexual antagonism were dependent on the founder number (Fig. 4A,B). When the founder number is two, equilibrium sex ratios in the three conditions are not significantly different, although we expect slight differences with a larger sample (*p* = 0.18; Tukey HSD test). The two-founder equilibria are closer to the paternal-only condition of the overlapping scenario than the maternal only condition. When founder numbers are low, less than six founders, sexual antagonism is greater in the overlapping simulation, consistent with the proposed hypothesis (Fig. 4B). However, when founder numbers are greater than six, we see no significant difference in the rate of sexual antagonism evolution for overlapping and discrete simulations (*p* < 10^−15^, *p* < 10^−15^, *p* < 10^−9^, *p* < 10^−5^ *p* < 0.67; t-test for two, three, four, five, and six founders respectively). For the two-founder scenario sexual antagonism in the discrete simulation in the final generation is significantly different than zero although the effect is small (*p* < 10^−10^, 95% CI [0.375 0.540]; t-test).

**Figure 4.**
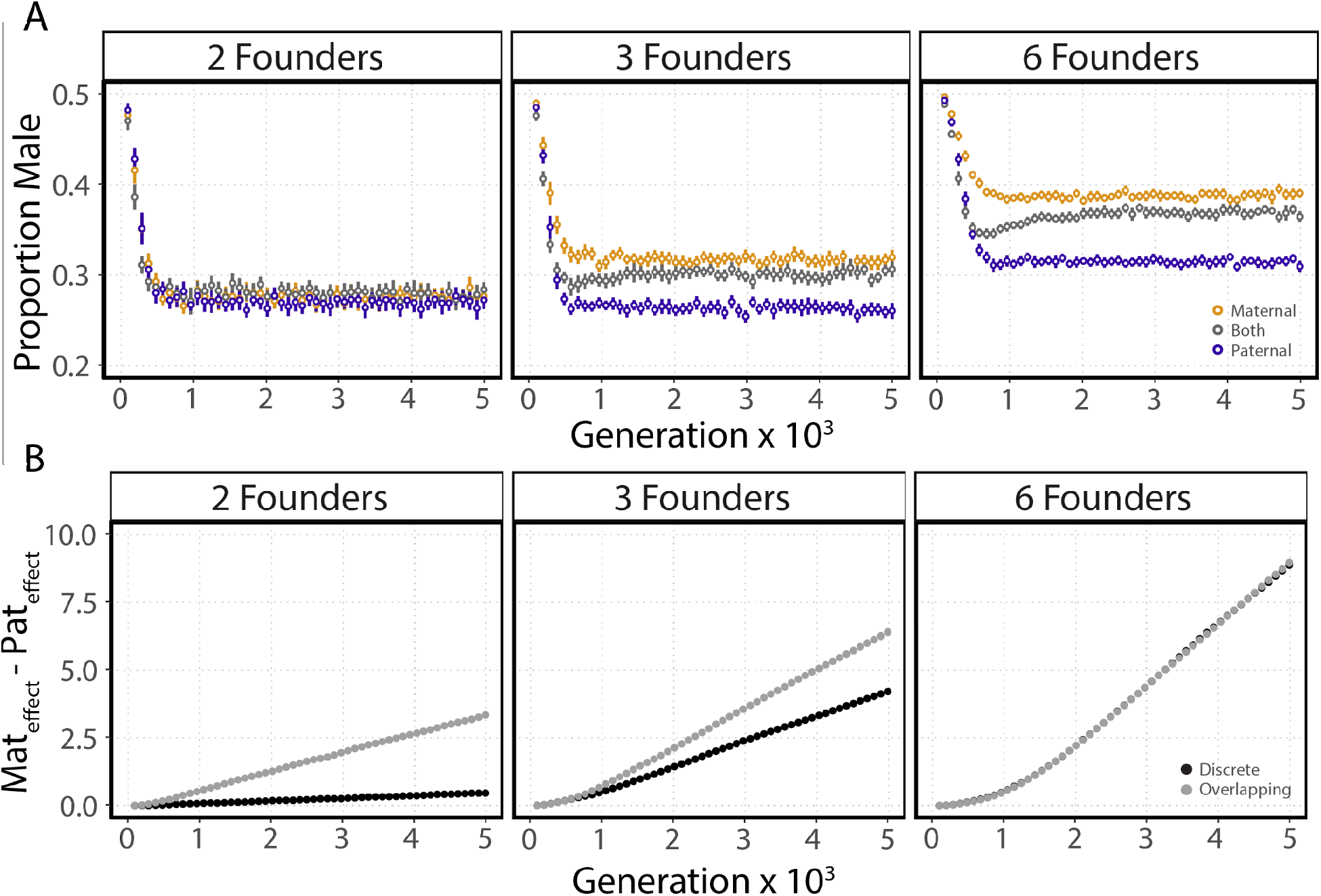
Overlapping generations affect the evolution of sexual antagonism when founder number is low. **(A)** Points represent mean population sex ratios measured in the first generation after patch founding at 96-generation intervals (bars represent the 95% CI of 25 simulations; brood size = 10, generations before migration = 3). **(B)** Points represent the mean of the difference in maternal genetic effects and paternal genetic effects measured in the first generation after patch founding at 96 generation intervals in the both maternal and paternal loci condition (bars represent the 95% CI of 25 simulations; brood size = 10, generations before migration = 3).

### Within- and among-patch variance differ between inheritance modes

Evolution of sex ratios in structured populations can be understood in terms of the variance of sex-ratio genotypes or strategies among patches (Aviles, 1993; Bulmer & Taylor, 1980b; Colwell, 1981; Frank, 1986; Hamilton, 1979; Wilson & Colwell, 1981). Patches with more female-biased sex ratios have a population growth advantage and will be overrepresented during migrant selection. When the variance among patches in migrant production is zero all patches will be equally represented in the migrant population and frequency-dependent forces will drive sex ratios to equality (Charnov, 1982). Across a range of founder numbers, we calculated the proportion of the variance in maternal or paternal genetic effects contained among patches out of the total variance of those effects in the founding generation. Maternal effects were calculated for the female population and paternal effects for the male population, although both sexes carry the other*’*s unexpressed loci. This value is equivalent to the population genetic estimator QST, the fixation index of a quantitative trait. Hamilton derived the evolutionarily stable strategy as a function of the within-patch variance over the total variance, which is the complement of this value (Hamilton, 1979).

Paternal genetic effect Q_ST_ is larger than maternal effect Q_ST_ for conditions with only maternal or paternal loci (Fig. 5A-D) and for simulations with both maternal and paternal loci (Fig. 5E-F). This is consistent with the hypothesis that Q_ST_ is driving equilibrium differences between maternal- and paternal-only simulations and sexual antagonism. However, it does not totally explain the phenomenon. Q_ST_ in the two-founder scenario is the same in overlapping and discrete simulations while the sex ratio in discrete simulations is less biased (Fig. 5B,D). This likely represents constraint at the first generation of offspring production on a patch. In the overlapping scenario a single male founder can assure mating for the second generation and is therefore free to evolve a more biased sex ratio. This constraint is likely not limited to just two founders. Sex ratios evolved under maternal-only conditions are also overall more biased with discrete generations (Fig. 5B,D). In both overlapping and discrete conditions, Q_ST_ explains the majority of the variation in sex ratios (overlapping, R^2^ = 0.8864; discrete, R^2^ = 0.8576). For conditions with both maternal and paternal loci, Q_ST_ is larger for paternal effects and smaller for maternal effects (Fig. 5E,F). Sex ratios across all simulation conditions and founder numbers evolved less-biased sex ratios than predicted by Hamilton*’*s model although our simulations do not match all of the assumptions of the model.

**Figure 5.**
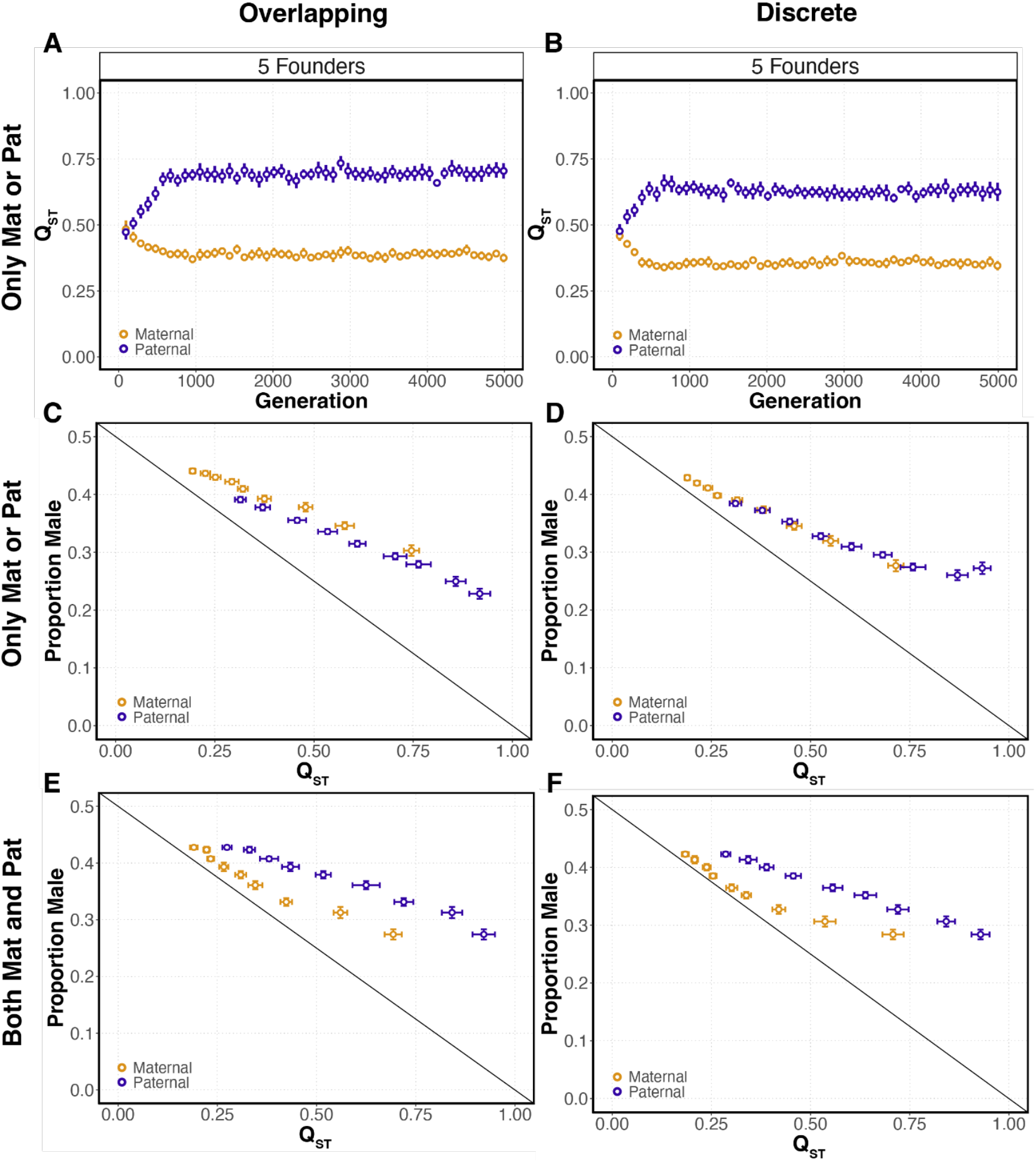
Q_ST_ is different for maternal and paternal genetic effects and predicts evolved sex ratios. **(A-B)** Points represent the Q_ST_ of maternal and paternal genetic effects measured in the founding generation at 96-generation intervals for simulations with overlapping and discrete generations. These are simulations with maternal or paternal loci only (bars represent the 95% CI of 25 simulations; brood size = 10, generations before migration = 3). **(C-D)** Points represent mean Q_ST_ and mean of the evolved sex ratio for each founder number 2-10. These values are measured in the founding generation at generation 4989 for simulations with maternal or paternal effects only (bars represent the 95% CI of 25 simulations; brood size = 10, generations before migration = 3). **(E-F)** Points represent mean Q_ST_ and mean of the evolved sex ratio for each founder number 2-10. These values are measured in the founding generation at generation 4989 for simulations with both maternal and paternal effects over a range of founders (bars represent the 95% CI of 25 simulations; brood size = 10, generations before migration = 3). Simulations have either overlapping (A, C, E) or discrete (B, D, F) generations. The diagonal lines represent the evolutionarily stable strategy for the relationship between Q_ST_ and the sex ratio from Hamilton*’*s 1979 group selection model.

### Effects of multiple generations and mating before and after dispersal

Metapopulations and the population biology of the species that live in them vary considerably in their details. This of course can affect both the sex-ratio equilibrium of populations and the rate at which they reach that equilibrium. Our simulations thus far have used three generations within a patch, with mating occurring after dispersal, in the next founding generation. We explored the effects of the number of generations in a patch, as well as the effects of other breeding schemes, including mating within a patch before dispersal (i.e., mated females disperse) and mating in a panmictic phase after dispersal from patches but prior to colonization of new patches. These population structures more closely resemble the LMC and haystack or structured demes models (Bulmer & Taylor, 1980b; Hamilton, 1967; Wilson & Colwell, 1981).

As predicted by the haystack model, sex ratios became more female biased as a function of the number of generations in a patch for all three models (Fig. 6). As expected, sex ratios do not evolve a bias with single-generation patches in either of the models with mating after dispersal, as they are equivalent to constant panmictic conditions. Overall, the inheritance mode did affect the evolved sex ratio for all models (*p* < 10^−4^, *p* < 10^−11^ and *p* < 10^−15^; deviance from linear regression for mating after colonization, mating in pool and mating before dispersal). In addition, we observed the evolution of sexual antagonism for each of the three models (Supp. Fig. 1).

**Figure 6.**
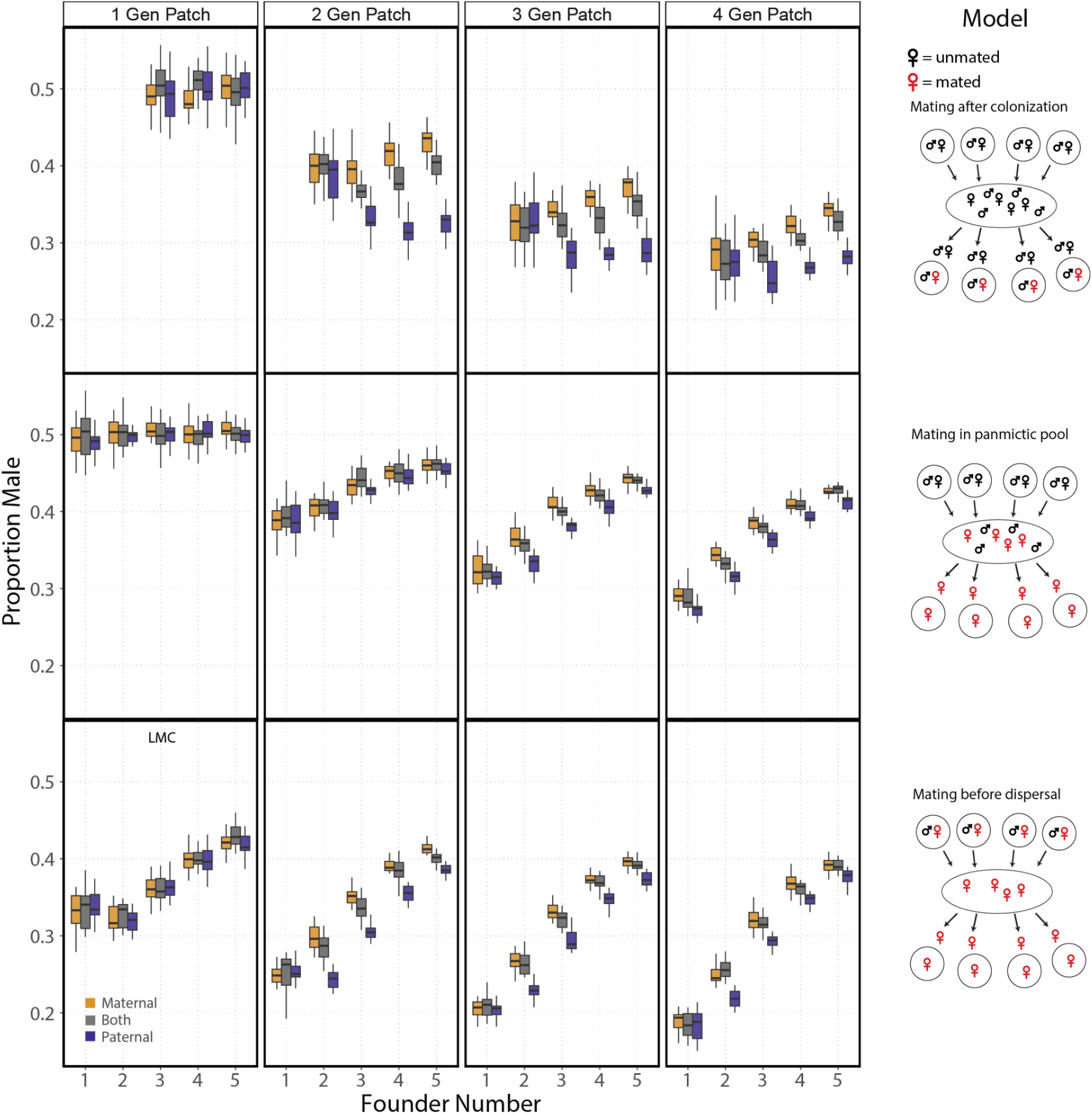
Comparing models of metapopulation dispersal and generations within a patch on equilibrium sex ratios for each inheritance mode and with discrete generations. Box plots for the sex ratio in generations 2588-2592 depending on the number of generations in a patch and the model (each condition was run 20 times; brood size = 5). A minimum of two founders is required for the mating-after-colonization model. Additionally, all populations in the two-founder single-generation patch experienced extinction. The one-generation patch with mating before dispersal is equivalent to LMC. Note, sex ratio under the one-founder one-generation simulation is constrained by brood size.

For the mating-before-dispersal model there was no difference in the evolved sex ratios between the inheritance modes with a one-generation patch, which is equivalent to the canonical LMC model (*p* = 0.90; deviance from linear regression). Similarly, we saw no difference in the evolved sex ratios between inheritance modes for the single-founder scenario across all numbers of generations in a patch for the mating-before-dispersal model (*p* = 0.96; deviance from linear regression). This was also true for mating after colonization for a founder number of two (*p* = 0.78; deviance from linear regression, see Fig. 4 for comparison). However, we did see a marginally significant difference when looking at the mating-in-pool model (*p* = 0.03; deviance from linear regression). These tests may be limited by simulation sample size, as we do observe the evolution of sexual antagonism for one and two founders in the mating-after-colonization and mating-in-pool models (Sup. Fig. 1).

## Discussion

We show for the first time that maternal or paternal autosomal control of the sex ratio leads to dramatically different equilibrium sex ratios in structured populations. Sex ratios that evolve under paternal control are more female biased. This effect is dependent on the population subdivision, where fewer founding individuals leads to both more biased sex ratios and a greater difference in the paternal and maternal equilibria. These results apply to species where autosomal loci that act paternally or maternally are sufficient to influence offspring sex probabilities.

This difference in evolved sex-ratio equilibria can be largely explained by Q_ST_ of the founding generation. This value represents the ratio of between-patch variance in maternal or paternal genetic effects over the total variance of those maternal or paternal genetic effects. Paternal Q_ST_ is higher than maternal Q_ST_ for maternal- and paternal-only conditions. This effect is dependent on the number of founders. Sex ratio is linearly correlated with Q_ST_ over a range of founders and Q_ST_ explains the majority of the variance in sex ratio equilibrium. This agrees with previous theory by Hamilton who used the Price equation to calculate the evolutionarily stable strategy as a function of the within-group variance over the total (Frank, 1986; Hamilton, 1979). So why might populations evolving under maternal and paternal control have different Q_ST_? The intuitive explanation relies on differences in sampling error for founding male and female genotypes. Only when there is variance in migrant production between patches is selection able to increase the frequency of female-biasing alleles (Colwell, 1981; Wilson & Colwell, 1981). Without this variance, within-patch frequency-dependent selection drives sex ratios to equality (Charnov, 1982). The mechanism producing this among-patch variation is sampling error in the migrant selection process, which causes sex-ratio genotypes to be distributed unevenly among patches. Sampling error increases as founder number decreases, leading to more biased sex ratios. As a consequence of the selection for a female-biased sex ratio, the population of males is smaller than the population of females. It follows that the average patch will have fewer male founders than female founders. Sampling error for males and their paternal genetic effects will consequently be higher, leading to greater variance among patches in population growth and eventually migrant production. Another way to describe this is that paternal-effect loci experience selection under a smaller “effective founder number” than maternal-effect loci given the same census founder number.

A second new result is that sexual antagonism evolves when maternal and paternal sex-ratio loci are acting in the same population. In this case, paternal loci that are female biasing are fixed continuously while maternal loci that are male biasing are fixed. The rate of sexual antagonism evolution is dependent on the number of founding individuals and peaks at six founders. These results can also be explained by Q_ST_; paternal-genetic-effect Q_ST_ is larger than maternal in the founding generation within the same simulations. These values stabilize sex ratios at an intermediate value closer to the maternal-only equilibrium sex ratio. One possible explanation for the observed peak at six founders is constraint, where having increasingly extreme genetic effects is detrimental to patch growth. Founding females with biases that exceed their mate*’*s produce a brood composed of mostly or only male offspring. Having larger numbers of founders lowers the probability of having a mating of incompatible genotypes leading to severely reduced patch output. This would reduce the rate of sexual antagonism evolution as genotypes with effects much larger than the population average would be selected against.

Generally, sex-ratio models have described populations with discrete generations (West, 2009). Our simulations are motivated by *Caenorhabditis* populations, where overlapping generations are the norm. Simulations with overlapping and discrete generations reveal several interesting results. When generations are overlapping, mating in subsequent generations is nearly guaranteed, as males can mate with daughters. In our simulations, we see constraints over the evolution of the sex ratio in discrete-generation simulations for paternally controlled sex. This effect is most apparent in the two-founder case, where the evolved sex ratio is the same as in the four-founder case rather than more extreme. This is likely the result of mating assurance, where male founders evolve less biased sex ratios under discrete generations because an all-female brood would end that patch*’*s lineage. Of more interest is that populations evolving under maternal control have more biased sex ratios with discrete generations. The reason for the shift is unlikely to be a difference in Q_ST_, which is nearly the same in overlapping vs. discrete simulations. Instead, this shift is possibly due to within-patch effects not captured by the Q_ST_ statistic.

The conclusions made here may seem at odds with sex ratio models that use the logic of inclusive fitness to calculate the unbeatable strategy. One common assumption is that founding mothers must have the same inclusive fitness as their mates and therefore it is only necessary to consider one parent, usually the mother. Our results don*’*t dispute this assumption. Rather, we conclude that paternal loci and maternal loci compete not with each other but against other loci of the same kind. The distribution of maternal and paternal effects among patches are uncorrelated as founders are distributed randomly each cycle. These distributions are summed each cycle such that over many cycles selection is acting on maternal and paternal loci independently. Using the logic of inclusive fitness we can imagine that founding mothers and fathers compete within a patch not with each other but with other founders of their same sex with some uncorrelated background effect from their mate*’*s genotype. The result is an intermediate population sex ratio.

Another unique aspect of our model, taken from the natural history of *Caenorhabditis*, is that mating occurs after dispersal and colonization, in the founding generation. The model is similar to mating after dispersal in a pool (as in the haystack model of Bulmer and Taylor, 1980b), while effectively decreasing the number of founders per patch. To determine the generality of our results we performed simulations where mating occurs either locally before dispersal or after dispersal in a panmictic pool. In addition, we tested the effects of multiple generations in a patch. As predicted by previous theory, additional generations in a patch lead to more biased sex ratios for all three models, with mating before dispersal having the most female-biased sex ratios (Bulmer & Taylor, 1980b). In addition, sexual antagonism evolved under all three models. Differences in the evolved sex ratios and the rate of sexual antagonism evolution are smaller in the mating-in-pool and mating-before-dispersal models compared to the mating-after-colonization model. An explanation consistent with the proposed hypothesis is that mating in a pool and mating before dispersal decrease among-patch variance in paternal genetic effects. This is because males either mating locally or in a panmictic pool can mate with more founding mothers than is possible in the mating-after-colonization model. Importantly, this highlights that life history features that alter the composition of founders or their genotypes will affect the differences in evolved sex ratios for animals with maternal or paternal control.

A necessary consideration when detecting sexual antagonism is that sex ratio phenotypes may be differentially constrained in mothers or fathers. Sexually antagonistic loci have been shown to be under pleiotropic constraint (Mank et al., 2008; Pennell et al., 2016). The antagonism described in this paper is interesting in that the expressed phenotype is the sum of maternal and paternal effects, never expressed in isolation. This means that constraints may not be due to pleiotropy but rather the ability of mothers or fathers to control the sex ratio. For example, fathers may approach a limit in making sperm differentially competitive or mothers may reach a limit in making embryos selectively viable. These may have different costs for each sex or one sex may reach a limit before the other. If such limits exist, the sex-ratio equilibrium may default to the sex with the larger range of sex-ratio control.

There is currently no evidence for sexual antagonism over autosomal sex-ratio loci, though many empirical observations are unexplained by previous theory (Orzack, 2002; Greeff & Kjellberg 2022). A detailed examination of a wider variety of taxa is necessary to test the generality of sex-ratio theory and identify parameters and constraints not considered in classic models. Detecting maternal and paternal effects on sex ratio is challenging, requiring careful experimental design. One solution would be to mate animals in both directions to a reference strain, as in Shuker et al. (2006), to estimate parental-effect heritability. Heritability, however, does not address the direction of the genetic effects. Our simulations make explicit predictions that paternal effects bias more toward female offspring compared to maternal effects. To test these predictions, we can estimate the effects of specific loci using a quantitative genetics approach. Such data will reveal the direction in which loci are biasing the sex ratio and in which sex they are active.

## Acknowledgments

This work was supported by grants GM141906 and HG013015. We thank Jon Wilkins, Peter Zee, and members of the Rockman lab for valuable discussion and advice, and NYU HPC staff for support.

---

**Supplementary Figure 1.**
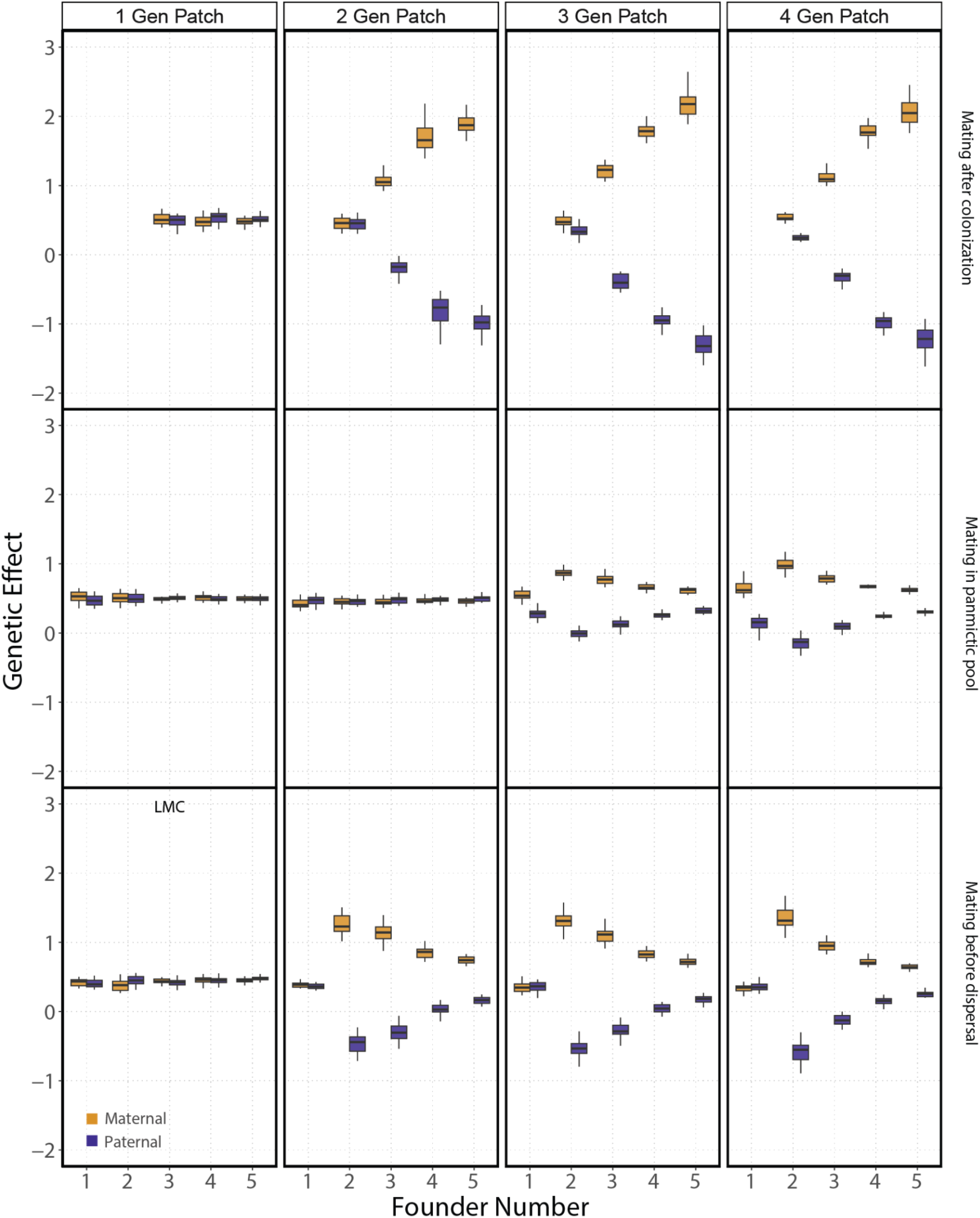
Comparing models of metapopulation dispersal and generations within a patch on evolved genetic effects for simulations with only maternal or paternal inheritance modes with discrete generations. Box plots for the genetic effects in generations 2588-2592 depending on the number of generations in a stack and the model (each condition was run 20 times; brood size = 5). A minimum of two founders is required for the mating after colonization model, additionally, all populations in the two-founder single-generation patch experienced extinction.

## Supplemental Text

Here we show single-locus recursion equations for a single patch, adapted from Wilson and Colwell*’*s (1981) haplodiploid model to the autosomal parental-effect diploid case with arbitrary dominance. These equations track the number of individuals of each sex and genotype in a growing patch. Terms that differ between the maternal- and paternal-effect models are highlighted in **red**.

|  |  |
| --- | --- |
| $N_f$ | Number of females |
| $N_m$ | Number of males |
| $N$ | Number of individuals ($N_f + N_m$) |
| $p_{f11}, p_{f12}, p_{f22}$ | Genotype frequencies among females (sum to 1) |
| $p_{m11}, p_{m12}, p_{m22}$ | Genotype frequencies among males (sum to 1) |
| $p_{f1}, p_{f2}$ | Allele frequencies among females (sum to 1) |
| $p_{m1}, p_{m2}$ | Allele frequencies among males (sum to 1) |
| $E$ | Number of offspring per female |
| $r$ | Sex ratio ($N_m/N$) in broods of $_{11}$ homozygotes of the ratio-controlling parent |
| $s$ | Shift of sex ratio in broods of $_{22}$ homozygotes of the ratio-controlling parent |
| $h$ | Dominance coefficient (i.e., ratios of $_{11}$, $_{12}$, and $_{22}$ are $r, r + hs, r + s$) |

Maternal-effect Recursion Equations

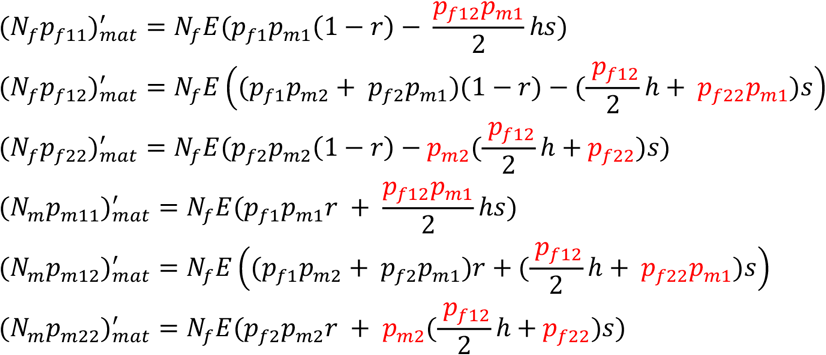

Paternal-effect Recursion Equations

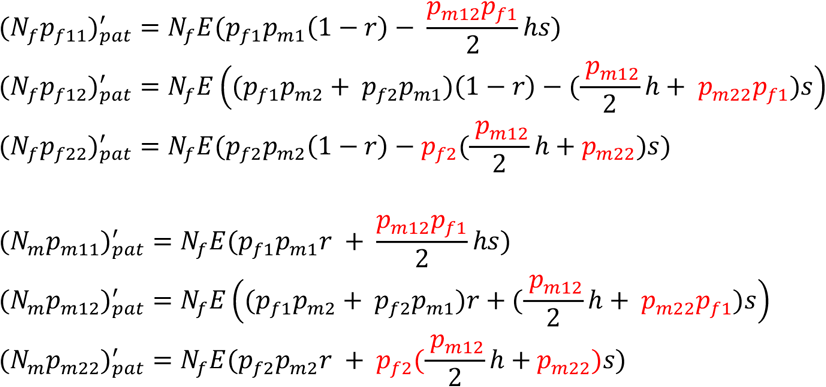

The within-patch differences between maternal and paternal control of the sex ratio can be illustrated with a simple numerical example involving a patch with three male and three female founders. In each patch, an allele that shifts offspring sex ratio from 0.5 to 0.35 to 0.2 with each additional allele in the parental genotype starts at frequency 0.5 in the founding generation. Each female has 10 offspring.

This example uses the following model parameters:

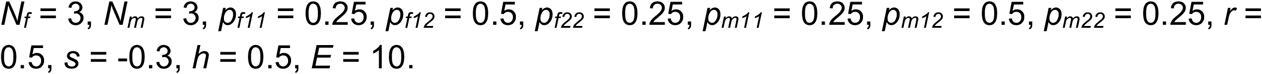

Paternal Control

| Generation | Individual Count | Sex Ratio | Allele Frequency |
| --- | --- | --- | --- |
| 1 | 6 | 0.5 | 0.500 |
| 2 | 30 | 0.350 | 0.500 |
| 3 | 195 | 0.366 | 0.488 |
| 4 | 1236.2 | 0.369 | 0.477 |
| 5 | 7803.6 | 0.371 | 0.467 |

Maternal Control

| Generation | Individual Count | Sex Ratio | Allele Frequency |
| --- | --- | --- | --- |
| 1 | 6 | 0.5 | 0.500 |
| 2 | 30 | 0.350 | 0.500 |
| 3 | 195 | 0.341 | 0.488 |
| 4 | 1284.4 | 0.345 | 0.474 |
| 5 | 8409.8 | 0.349 | 0.462 |

