## Supplementary material for "Sexual antagonism evolves when autosomes influence offspring sex ratio": FiguresAndAnalysis.R

```
#Supplementary file to reproduce analysis and figures in, "Sexual antagonism evolves when autosomes influence offspring sex ratio " by Solomon Sloat and Matthew Rockman.

#load libraries
library(ggplot2)
library(tidyverse)
library(gridExtra)
library(grid)
library(plyr)
library(Rmisc)
library(ggpubr)
library(readxl)

##############################
### Load data and reformat ###
##############################

#See key in "Supplementary_data.xlsx" for description of columns and simulation parameters. 

#imort data for overlapping generations br=10
SR_Mode <- read_excel("Supplementary_data.xlsx", sheet = "Supplemental_Data_1")
SR_Mode <- as_tibble(SR_Mode)
SR_Mode$mode[SR_Mode$mode =='1'] = 'F'
SR_Mode$mode[SR_Mode$mode =='2'] = 'M'
SR_Mode$mode[SR_Mode$mode =='3'] = 'F&M'
SR_Mode$mode <- as.factor(SR_Mode$mode)
SR_Mode$nFounders <- as.factor(SR_Mode$nFounders)
SR_Mode$mode <- factor(SR_Mode$mode, levels = c("F", "F&M", "M"))
SR_Mode$nFounders_char <- paste(SR_Mode$nFounders,'Founders', sep = ' ')
SR_Mode$nFounders_char <- factor(SR_Mode$nFounders_char, levels = c("2 Founders", "3 Founders", "4 Founders",
                                                          "5 Founders", "6 Founders", "7 Founders",
                                                          "8 Founders", "9 Founders", "10 Founders"))

#import data for overlapping generations br=3 founder number 2-30
SR_Mode30 <-read_excel("Supplementary_data.xlsx", sheet = "Supplemental_Data_2")
SR_Mode30 <-as_tibble(SR_Mode30)
SR_Mode30$mode[SR_Mode30$mode =='1'] = 'F'
SR_Mode30$mode[SR_Mode30$mode =='2'] = 'M'
SR_Mode30$mode[SR_Mode30$mode =='3'] = 'F&M'
SR_Mode30$mode <- as.factor(SR_Mode30$mode)

# import data for descrete generations
SR_Disc <- read_excel("Supplementary_data.xlsx", sheet = "Supplemental_Data_3")
SR_Disc <- as_tibble(SR_Disc)
SR_Disc$mode[SR_Disc$mode =='1'] = 'F'
SR_Disc$mode[SR_Disc$mode =='2'] = 'M'
SR_Disc$mode[SR_Disc$mode =='3'] = 'F&M'
SR_Disc$mode <- as.factor(SR_Disc$mode)
SR_Disc$nFounders <- as.factor(SR_Disc$nFounders)
SR_Disc$mode <- factor(SR_Disc$mode, levels = c("F", "F&M", "M"))
SR_Disc$nFounders_char <- paste(SR_Disc$nFounders,'Founders',sep = ' ')
SR_Disc$nFounders_char <- factor(SR_Disc$nFounders_char, levels = c("2 Founders", "3 Founders", "4 Founders",
                                                          "5 Founders", "6 Founders", "7 Founders",
                                                          "8 Founders", "9 Founders", "10 Founders"))

#import data for mating after dispersal in a pool
SR_AD <- read_excel("Supplementary_data.xlsx", sheet = "Supplemental_Data_5")
SR_AD <-as_tibble(SR_AD)
SR_AD$nFounders <- as.factor(as.character(SR_AD$nFounders))
SR_AD$G<- paste(SR_AD$G,'Gen Patch',sep = ' ')
SR_AD$G <- as.factor(SR_AD$G)
SR_AD$mode = factor(SR_AD$mode,
                    levels = c("1", "3", "2"))

#import data for mating before dispersal locally
SR_BD <- read_excel("Supplementary_data.xlsx", sheet = "Supplemental_Data_6")
SR_BD <-as_tibble(SR_BD)
SR_BD$nFounders <- as.factor(as.character(SR_BD$nFounders))
SR_BD$G<- paste(SR_BD$G,'Gen Patch',sep = ' ')
SR_BD$G <- as.factor(SR_BD$G)
SR_BD$mode = factor(SR_BD$mode,
                    levels = c("1", "3", "2"))

#import data for mating after dispersal and colonization (females and males are founders)
SR_MAC <- read_excel("Supplementary_data.xlsx", sheet = "Supplemental_Data_4")
SR_MAC <-as_tibble(SR_MAC)
SR_MAC$nFounders <- as.factor(as.character(SR_MAC$nFounders))
SR_MAC$G<- paste(SR_MAC$G,'Gen Patch',sep = ' ')
SR_MAC$G <- as.factor(SR_MAC$G)
SR_MAC$mode = factor(SR_MAC$mode,
                     levels = c("1", "3", "2"))

###################################################################
### Does the mode of inheritance affect equilibrium sex ratios? ###
###################################################################

#calculate summary statistics
SR_Mode_SE <- summarySE(SR_Mode, measurevar="SexRatio", groupvars=c("generation","nFounders_char", "mode"))

#plot sex ratio by generation 
p1 <- ggplot(SR_Mode_SE, aes(x=generation, y=SexRatio, color=mode, shape=mode,fill='white')) + 
  geom_errorbar(aes(ymin=SexRatio-ci, ymax=SexRatio+ci),width=0,size=1) +
  geom_point(aes(stroke=1),size=1) +
  scale_x_continuous( limits=c(0,5000)) +
  theme_classic() +
  ylab("Proportion Male") +
  xlab("Generation") +
  ylim(0.2,0.5) +
  theme(axis.text=element_text(size=20),
        axis.title=element_text(size=25,face="bold"), strip.background =element_rect(fill="white"), strip.text = element_text(size = 25),
        legend.position = "none", panel.border = element_rect(colour = "black", fill=NA, linewidth=2), panel.grid.major = element_line(color = 'lightgrey', linetype = 'dotted')) + 
  scale_shape_manual(values = c(21, 21, 21)) +
  scale_color_manual(values = c('#D9911A', '#666666', '#3305A3')) +
  scale_linetype_manual(values =c(1,1,1)) +
  scale_fill_manual(values ='white')

wrapped_p1 <- p1 + facet_wrap(~nFounders_char,nrow =3)

wrapped_p1

#plot sex ratio for founder number 5 (Figure 2A) 
p2 <- ggplot(filter(SR_Mode_SE, nFounders_char == "5 Founders"), aes(x=generation, y=SexRatio, color=mode, shape=mode,fill='white')) + 
  geom_errorbar(aes(ymin=SexRatio-ci, ymax=SexRatio+ci),width=0,size=1.5) +
  geom_point(aes(stroke=1.75),size=2) +
  scale_x_continuous( limits=c(0,5000)) +
  theme_classic() +
  ylab("Proportion Male") +
  xlab("Generation") +
  ylim(0.25,0.5) +
  theme(axis.text=element_text(size=20),
        axis.title=element_text(size=25,face="bold"), strip.background =element_rect(fill="white"), strip.text = element_text(size = 25),
        legend.position = "none", panel.border = element_rect(colour = "black", fill=NA, linewidth=2), panel.grid.major = element_line(color = 'lightgrey', linetype = 'dotted')) + 
  scale_shape_manual(values = c(21, 21, 21)) +
  scale_color_manual(values = c('#D9911A', '#666666', '#3305A3')) +
  scale_linetype_manual(values =c(1,1,1)) +
  scale_fill_manual(values ='white')

wrapped_p2 <- p2 + facet_wrap(~nFounders_char,nrow =3)

wrapped_p2


#Does mode affect the evolved sex ratio?

lm1_mode=lm(data = SR_Mode, formula = SexRatio ~ mode + nFounders + generation)
lm2_mode=lm(data = SR_Mode, formula = SexRatio ~ nFounders + generation)

anova(lm1_mode,lm2_mode, test = "Chisq")

"""
Analysis of Variance Table

Model 1: firstGenSexRatio ~ mode + nFounders + generation
Model 2: firstGenSexRatio ~ nFounders + generation
  Res.Df     RSS Df Sum of Sq  Pr(>Chi)    
1  70188  49.914                           
2  70190 126.708 -2   -76.794 < 2.2e-16 ***
---
Signif. codes:  0 ‘***’ 0.001 ‘**’ 0.01 ‘*’ 0.05 ‘.’ 0.1 ‘ ’ 1
"""

# Plot sex ratio at equilibrium for founder number 2-30, brood size = 3. (Figure 2B)

SR_Mode30_SE <- summarySE(SR_Mode30, measurevar="SexRatio", groupvars=c("generation","nFounders", "mode"))

p3<- ggplot(filter(SR_Mode30_SE, generation == 4989), mapping=aes(nFounders,SexRatio,color=mode)) + 
  geom_errorbar(aes(ymin=SexRatio-ci, ymax=SexRatio+ci),width=0,size=1.5) +
  stat_summary(fun.y=mean, geom="point", stroke=2, size=2, fill='white', aes(shape=mode,color=mode)) +
  scale_x_continuous(breaks=c(2,4,6,8,10,15,20,25,30)) +
  ylim(0.2,0.5) +
  scale_shape_manual(values = c(21, 21, 21)) +
  scale_color_manual(values = c('#D9911A','#666666', '#3305A3')) +
  ylab("Proportion Male") +
  xlab("Founder Number") +
  theme_classic() +
  theme(axis.text=element_text(size=20),
        axis.title=element_text(size=25,face="bold"), strip.text = element_text(size = 18),
        legend.position = "none", panel.border = element_rect(colour = "black", fill=NA, linewidth=2), panel.grid.major = element_line(color = 'lightgrey', linetype = 'dotted'))

p3

# Does brood size constrain the sex ratio?

SR_Mode30$broodsize <- "3"
SR_Mode$broodsize <- "10"

br10_br3 <- rbind(SR_Mode30,SR_Mode[,-10])
br10_br3_filt <- filter(br10_br3, nFounders < 11, generation == 4989)

lm1_br10_br3=lm(data = br10_br3_filt, formula = SexRatio ~ mode + nFounders + broodsize)
lm2_br10_br3=lm(data = br10_br3_filt, formula = SexRatio ~  mode + nFounders)

anova(lm1_br10_br2, lm2_br10_br2, test = "Chisq")

"""
Analysis of Variance Table

Model 1: firstGenSexRatio ~ mode + nFounders + generation + broodsize
Model 2: firstGenSexRatio ~ mode + nFounders + generation
  Res.Df    RSS Df Sum of Sq  Pr(>Chi)    
1 140394 135.48                           
2 140395 143.42 -1    -7.939 < 2.2e-16 ***
---
Signif. codes:  0 ‘***’ 0.001 ‘**’ 0.01 ‘*’ 0.05 ‘.’ 0.1 ‘ ’ 1
"""

#############################################################
### Are maternal and paternal loci sexually antagonistic? ###
#############################################################


SR_Mode %>% pivot_longer(cols=c('SexRatio', 'matGE', 'patGE'),
                             names_to='names',
                             values_to='values') -> GE_Mode

GE_Mode_SE <- summarySE(filter(GE_Mode, mode=="F&M"), measurevar="values", groupvars=c("generation","nFounders_char","names"))
GE_Mode_SE$names <- factor(GE_Mode_SE$names)
levels(GE_Mode_SE$names) <- c("matGE","SexRatio", "patGE")

#plot genetic effects and sex ratio by generation
p4 <- ggplot(GE_Mode_SE, aes(x=generation, y=values, color=names, shape=names, fill='white')) + 
  geom_errorbar(aes(ymin=values-ci, ymax=values+ci),width=0,size=1) +
  geom_point(aes(stroke=1),size=1) +
  scale_x_continuous(limits=c(0,5000)) +
  theme_classic() +
  ylab("Genetic Effect") +
  xlab("Generation") +
  #ylim(-.5,1.25) +
  theme(axis.text=element_text(size=20),
        axis.title=element_text(size=25,face="bold"), strip.background =element_rect(fill="white"), strip.text = element_text(size = 25),
        legend.position = "none", panel.border = element_rect(colour = "black", fill=NA, linewidth=2), panel.grid.major = element_line(color = 'lightgrey', linetype = 'dotted')) + 
  scale_shape_manual( values = c(23, 23, 21)) +
  scale_color_manual(values = c('#D9911A','#3305A3','#666666')) +
  scale_fill_manual(values ='white')

wrapped_p4 <- p4 + facet_wrap(~nFounders_char)
wrapped_p4

#plot genetic effects and sex ratio by generation (Figure 3A)
p5 <- ggplot(filter(GE_Mode_SE, nFounders_char == "5 Founders"), aes(x=generation, y=values, color=names, shape=names, fill='white')) + 
  geom_errorbar(aes(ymin=values-ci, ymax=values+ci),width=0,size=2) +
  geom_point(aes(stroke=1.75),size=2) +
  scale_x_continuous(limits=c(0,5000)) +
  scale_y_continuous(limits=c(-7,7), sec.axis = sec_axis(~ . * 1, name = "Proportion Male")) +
  theme_classic() +
  ylab("Genetic Effect") +
  xlab("Generation") +
  #ylim(-7,7) +
  theme(axis.text=element_text(size=20),
        axis.title=element_text(size=25,face="bold"), strip.background =element_rect(fill="white"), strip.text = element_text(size = 25),
        legend.position = "none", panel.border = element_rect(colour = "black", fill=NA, linewidth=2), panel.grid.major = element_line(color = 'lightgrey', linetype = 'dotted')) + 
  scale_shape_manual(values = c(23, 23, 21)) +
  scale_color_manual(values = c('#D9911A','#3305A3','#666666')) +
  scale_fill_manual(values ='white')

wrapped_p5 <- p5 + facet_wrap(~nFounders_char)
wrapped_p5


# plot genetic effects by founder in the final generation (Figure 3B)

SR_Mode30 %>% pivot_longer(cols=c('SexRatio', 'matGE', 'patGE'),
                             names_to='names',
                             values_to='values') -> GE_Mode30

GE_Mode30_SE <- summarySE(filter(GE_Mode30, mode=="F&M"), measurevar="values", groupvars=c("generation","nFounders","names"))
GE_Mode30_SE$names <- factor(GE_Mode30_SE$names)
levels(GE_Mode30_SE$names) <- c("matGE","SexRatio", "patGE")

p6<- ggplot(filter(GE_Mode30_SE, generation == max(GE_Mode30_SE$generation)), mapping=aes(nFounders,values,color=names, fill = 'white')) + 
  geom_errorbar(aes(ymin=values-ci, ymax=values+ci),width=0,size=1.5) +
  stat_summary(fun.y=mean, geom="point", stroke=2, size=2, fill='white', aes(shape=names,color=names)) +
  scale_x_continuous(breaks=c(2,4,6,8,10,15,20,25,30)) +
  scale_y_continuous(limits=c(-10,10), sec.axis = sec_axis(~ . * 1, name = "Proportion Male")) +
  scale_shape_manual(values = c(23, 23, 21)) +
  scale_color_manual(values = c('#D9911A','#3305A3','#666666')) +
  ylab("Genetic Effect") +
  xlab("Founder Number") +
  theme_classic() +
  theme(axis.text=element_text(size=20),
        axis.title=element_text(size=25,face="bold"), strip.text = element_text(size = 18),
        legend.position = "none", panel.border = element_rect(colour = "black", fill=NA, linewidth=2), panel.grid.major = element_line(color = 'lightgrey', linetype = 'dotted'))

p6

ggsave(plot = p3,filename = 'GE_byFounder_br3_Fdrs2to30_3genStack_OG_032423_v2.pdf', height = 6, width = 8, dpi = 320)


###############################################################################
### Does overlapping vs discrete generations affect equilibrium sex ratios? ###
###############################################################################


#calculate summary statistics for sex ratios for overlapping and discrete generations
SR_Disc_SE <- summarySE(SR_Disc, measurevar="SexRatio", groupvars=c("generation","nFounders_char", "mode"))

#plot sex ratios by generation for overlapping and discrete generations (Figure 4A)
p6 <- ggplot(filter(SR_Disc_SE, nFounders_char != '4 Founders', nFounders_char != '5 Founders', nFounders_char != '7 Founders', nFounders_char != '8 Founders', nFounders_char != '9 Founders', nFounders_char != '10 Founders'), aes(x=generation, y=SexRatio, color=mode, shape=mode,fill='white')) + 
  geom_errorbar(aes(ymin=SexRatio-ci, ymax=SexRatio+ci),width=0,size=1) +
  geom_point(aes(stroke=1),size=1) +
  scale_x_continuous( limits=c(0,5000)) +
  theme_classic() +
  ylab("Proportion Male") +
  xlab("Generation") +
  ylim(0.2,0.5) +
  theme(axis.text=element_text(size=20),
        axis.title=element_text(size=25,face="bold"), strip.background =element_rect(fill="white"), strip.text = element_text(size = 25),
        legend.position = "none", panel.border = element_rect(colour = "black", fill=NA, linewidth=2), panel.grid.major = element_line(color = 'lightgrey', linetype = 'dotted')) + 
  scale_shape_manual(values = c(21, 21, 21)) +
  scale_color_manual(values = c('#D9911A', '#666666', '#3305A3')) +
  scale_linetype_manual(values =c(1,1,1)) +
  scale_fill_manual(values ='white')

wrapped_p6 <- p6 + facet_wrap(~nFounders_char,nrow = 1)
wrapped_p6


#generate summary statistics for maternal and paternal effects for overlapping and discrete generations
SR_Mode$isoverlapping <- "T"
SR_Disc$isoverlapping <- "F"
SR_OG_DISC <- rbind(SR_Disc, SR_Mode)
SR_OG_DISC$GE_prop <- SR_OG_DISC$matGE - SR_OG_DISC$patGE

filter(SR_OG_DISC, mode == "F&M") %>% pivot_longer(cols=c('SexRatio', 'matGE', 'patGE'),
                             names_to='names',
                             values_to='vales') -> GE_OG_DISC

GE_OG_DISC_SE <- summarySE(GE_OG_DISC, measurevar="GE_prop", groupvars=c("generation","nFounders_char","isoverlapping"))


#plot difference in maternal effects for overlapping and discrete generations (Figure 4B)
p7 <- ggplot(filter(GE_OG_DISC_SE, nFounders_char != '4 Founders', nFounders_char != '5 Founders', nFounders_char != '7 Founders', nFounders_char != '8 Founders', nFounders_char != '9 Founders', nFounders_char != '10 Founders'), aes(x=generation, y=GE_prop, color=isoverlapping, fill = "white")) + 
  geom_errorbar(aes(ymin=GE_prop-ci, ymax=GE_prop+ci),width=0,size=1) +
  geom_point(aes(stroke=1),size=1) +
  scale_x_continuous(limits=c(0,5000)) +
  theme_classic() +
  ylab("Maternal Effect - Paternal Effect") +
  xlab("Generation") +
  coord_cartesian(ylim=c(0, 10)) +
  theme(axis.text=element_text(size=20),
        axis.title=element_text(size=25,face="bold"), strip.background =element_rect(fill="white"), strip.text = element_text(size = 25),
        legend.position = "none", panel.border = element_rect(colour = "black", fill=NA, linewidth=2), panel.grid.major = element_line(color = 'lightgrey', linetype = 'dotted')) + 
  scale_shape_manual( values = c(21, 21, 21)) +
  scale_color_manual(values = c('#000000', '#A0A0A0','#666666')) +
  scale_fill_manual(values ='white')

wrapped_p7 <- p7 + facet_wrap(~nFounders_char)
wrapped_p7


#Are equilibrium sex ratios different in the two founder case?

two_founder_aov = aov(SexRatio ~ mode,
    data = filter(SR_Disc, generation == 4989, nFounders_char == "2 Founders"))
    
TukeyHSD(two_founder_aov)

"""
Tukey multiple comparisons of means
    95% family-wise confidence level

Fit: aov(formula = firstGenSexRatio ~ SA_QTL, data = filter(SRQTL_byQTL_OG, generation == 4989, nFounders == 2))

$SA_QTL
             diff          lwr         upr     p adj
F&M-F  0.00720940 -0.008267868 0.022686668 0.5081872
M-F   -0.00434188 -0.019819148 0.011135388 0.7808304
M-F&M -0.01155128 -0.027028548 0.003925988 0.1814928
"""

#Are the differences in sexual antagonism between overlapping and discrete conditions dependent on the number of founders?

diff_GE_2_founder_ttest = t.test(filter(SR_OG_DISC, generation == 4989, mode == "F&M", nFounders_char == "2 Founders", isoverlapping == "T")$GE_prop, 
                                   filter(SR_OG_DISC, generation == 4989, mode == "F&M", nFounders_char == "2 Founders", isoverlapping == "F")$GE_prop)
diff_GE_2_founder_ttest

"""
Welch Two Sample t-test

data:  filter(SR_OG_DISC, generation == 4989, mode == "F&M", nFounders == "2 Founders", isoverlapping == "T")$GE_prop and filter(SR_OG_DISC, generation == 4989, mode == "F&M", nFounders == "2 Founders", isoverlapping == "F")$GE_prop
t = 31.858, df = 35.099, p-value < 2.2e-16
alternative hypothesis: true difference in means is not equal to 0
95 percent confidence interval:
 2.697799 3.064993
sample estimates:
mean of x mean of y 
3.3394810 0.4580852 
"""

diff_GE_3_founder_ttest = t.test(filter(SR_OG_DISC, generation == 4989, mode == "F&M", nFounders_char == "3 Founders", isoverlapping == "T")$GE_prop, 
                                 filter(SR_OG_DISC, generation == 4989, mode == "F&M", nFounders_char == "3 Founders", isoverlapping == "F")$GE_prop)
diff_GE_3_founder_ttest

"""
Welch Two Sample t-test

data:  filter(SR_OG_DISC, generation == 4989, mode == "F&M", nFounders == "3 Founders", isoverlapping == "T")$GE_prop and filter(SR_OG_DISC, generation == 4989, mode == "F&M", nFounders == "3 Founders", isoverlapping == "F")$GE_prop
t = 13.285, df = 45.132, p-value < 2.2e-16
alternative hypothesis: true difference in means is not equal to 0
95 percent confidence interval:
 1.857998 2.521962
sample estimates:
mean of x mean of y 
 6.397462  4.207482 

"""

diff_GE_4_founder_ttest = t.test(filter(SR_OG_DISC, generation == 4989, mode == "F&M", nFounders_char == "4 Founders", isoverlapping == "T")$GE_prop, 
                                 filter(SR_OG_DISC, generation == 4989, mode == "F&M", nFounders_char == "4 Founders", isoverlapping == "F")$GE_prop)
diff_GE_4_founder_ttest

"""
Welch Two Sample t-test

data:  filter(SR_OG_DISC, generation == 4989, mode == "F&M", nFounders == "4 Founders", isoverlapping == "T")$GE_prop and filter(SR_OG_DISC, generation == 4989, mode == "F&M", nFounders == "4 Founders", isoverlapping == "F")$GE_prop
t = 8.1503, df = 47.918, p-value = 1.323e-10
alternative hypothesis: true difference in means is not equal to 0
95 percent confidence interval:
 1.080288 1.787885
sample estimates:
mean of x mean of y 
 8.389657  6.955570 

"""

diff_GE_5_founder_ttest = t.test(filter(SR_OG_DISC, generation == 4989, mode == "F&M", nFounders_char == "5 Founders", isoverlapping == "T")$GE_prop, 
                                 filter(SR_OG_DISC, generation == 4989, mode == "F&M", nFounders_char == "5 Founders", isoverlapping == "F")$GE_prop)
diff_GE_5_founder_ttest

"""
Welch Two Sample t-test

data:  filter(SR_OG_DISC, generation == 4989, mode == "F&M", nFounders == "5 Founders", isoverlapping == "T")$GE_prop and filter(SR_OG_DISC, generation == 4989, mode == "F&M", nFounders == "5 Founders", isoverlapping == "F")$GE_prop
t = 5.4823, df = 47.352, p-value = 1.589e-06
alternative hypothesis: true difference in means is not equal to 0
95 percent confidence interval:
 0.583209 1.259130
sample estimates:
mean of x mean of y 
 9.184994  8.263824 

"""

diff_GE_6_founder_ttest = t.test(filter(SR_OG_DISC, generation == 4989, mode == "F&M", nFounders_char == "6 Founders", isoverlapping == "T")$GE_prop, 
                                 filter(SR_OG_DISC, generation == 4989, mode == "F&M", nFounders_char == "6 Founders", isoverlapping == "F")$GE_prop)
diff_GE_6_founder_ttest

"""
Welch Two Sample t-test

data:  filter(SR_OG_DISC, generation == 4989, mode == "F&M", nFounders == "6 Founders", isoverlapping == "T")$GE_prop and filter(SR_OG_DISC, generation == 4989, mode == "F&M", nFounders == "6 Founders", isoverlapping == "F")$GE_prop
t = 0.42839, df = 47.524, p-value = 0.6703
alternative hypothesis: true difference in means is not equal to 0
95 percent confidence interval:
 -0.2680777  0.4131921
sample estimates:
mean of x mean of y 
 8.953766  8.881209 

"""
#Is sexual antagonism or the difference between paternal and maternal effects in the two founder discrete condition equal to zero?

GE_2_founder_null_zero_ttest = t.test(filter(SR_OG_DISC, generation == 4989, mode == "F&M", nFounders_char == "2 Founders", isoverlapping == "F")$GE_prop)

GE_2_founder_null_zero_ttest

"""
One Sample t-test

data:  filter(SR_OG_DISC, generation == 4989, mode == "F&M", nFounders == "2 Founders", isoverlapping == "F")$GE_prop
t = 11.415, df = 24, p-value = 3.492e-11
alternative hypothesis: true mean is not equal to 0
95 percent confidence interval:
 0.3752605 0.5409099
sample estimates:
mean of x 
0.4580852 

"""

###################################################################
### Is Qst different between maternal and paternal effect loci? ###
###################################################################


#This section is for overlapping generations.

SR_Mode %>% pivot_longer(cols=c('QstPat', 'QstMat'),
                             names_to='MatOrPat',
                             values_to='Qst') -> Qst_Mode

Qst_Mode_SE <- summarySE(Qst_Mode, measurevar="Qst", groupvars=c("generation","nFounders_char", "mode", "MatOrPat"))

c1 = Qst_Mode_SE[Qst_Mode_SE$mode == 'F' &  Qst_Mode_SE$MatOrPat == 'QstMat',]
c2 = Qst_Mode_SE[Qst_Mode_SE$mode == 'F&M' &  Qst_Mode_SE$MatOrPat == 'QstMat',]
c3 = Qst_Mode_SE[Qst_Mode_SE$mode == 'M' &  Qst_Mode_SE$MatOrPat == 'QstPat',]
c4 = Qst_Mode_SE[Qst_Mode_SE$mode == 'F&M' &  Qst_Mode_SE$MatOrPat == 'QstPat',]

Qst_Mode_SE_filt = rbind(c1,c3)
Qst_Mode_SE_filt$condition = factor(paste(Qst_Mode_SE_filt$MatOrPat, Qst_Mode_SE_filt$mode, sep = '_'))
Qst_Mode_SE_filt = na.omit(Qst_Mode_SE_filt)
Qst_Mode_SE_filt$condition = droplevels(Qst_Mode_SE_filt$condition)

#plot Qst by maternal and paternal effects in the maternal and paternal only conditions
p8 <- ggplot(Qst_Mode_SE_filt, aes(x=generation, y=Qst, color=condition, shape=condition,fill='white')) + 
  geom_errorbar(aes(ymin=Qst-ci, ymax=Qst+ci),width=0,size=1) +
  geom_point(aes(stroke=1),size=1) +
  scale_x_continuous( limits=c(0,5000)) +
  theme_classic() +
  ylab("Qst") +
  xlab("Generation") +
  ylim(0,1) +
  theme(axis.text=element_text(size=20),
        axis.title=element_text(size=25,face="bold"), strip.background =element_rect(fill="white"), strip.text = element_text(size = 25),
        legend.position = "none", panel.border = element_rect(colour = "black", fill=NA, linewidth=2), panel.grid.major = element_line(color = 'lightgrey', linetype = 'dotted')) + 
  scale_shape_manual(values = c(21, 21)) +
  scale_color_manual(values = c('#D9911A', '#3305A3')) +
  scale_fill_manual(values ='white')

wrapped_p8 <- p8 + facet_wrap(~nFounders_char)
wrapped_p8


#plot Qst by maternal and paternal effects with 5 founders for (Figure 5A)
p9 <- ggplot(filter(Qst_Mode_SE_filt, nFounders_char == "5 Founders"), aes(x=generation, y=Qst, color=condition, shape=condition,fill='white')) + 
  geom_errorbar(aes(ymin=Qst-ci, ymax=Qst+ci),width=0,size=1) +
  geom_point(aes(stroke=1),size=1) +
  scale_x_continuous( limits=c(0,5000)) +
  theme_classic() +
  ylab("Qst") +
  xlab("Generation") +
  ylim(0,1) +
  theme(axis.text=element_text(size=20),
        axis.title=element_text(size=25,face="bold"), strip.background =element_rect(fill="white"), strip.text = element_text(size = 25),
        legend.position = "none", panel.border = element_rect(colour = "black", fill=NA, linewidth=2), panel.grid.major = element_line(color = 'lightgrey', linetype = 'dotted')) + 
  scale_shape_manual(values = c(21, 21)) +
  scale_color_manual(values = c('#D9911A', '#3305A3')) +
  scale_fill_manual(values ='white')

wrapped_p9 <- p9 + facet_wrap(~nFounders_char)
wrapped_p9

# plot sex ratio by Qst and by founder in the final generation

Qst_Mode_SR_SE <- summarySE(Qst_Mode, measurevar="SexRatio", groupvars=c("generation","nFounders_char", "mode", "MatOrPat"))
Qst_Mode_Merge <- merge(Qst_Mode_SE,Qst_Mode_SR_SE,by=c("generation","nFounders_char","mode","MatOrPat"))

c1_Merge  = Qst_Mode_Merge[Qst_Mode_Merge$mode == 'F' &  Qst_Mode_Merge$MatOrPat == 'QstMat',]
c2_Merge  = Qst_Mode_Merge[Qst_Mode_Merge$mode == 'F&M' &  Qst_Mode_Merge$MatOrPat == 'QstMat',]
c3_Merge  = Qst_Mode_Merge[Qst_Mode_Merge$mode == 'M' &  Qst_Mode_Merge$MatOrPat == 'QstPat',]
c4_Merge  = Qst_Mode_Merge[Qst_Mode_Merge$mode == 'F&M' &  Qst_Mode_Merge$MatOrPat == 'QstPat',]

Qst_Mode_Merge_filt = rbind(c1_Merge,c3_Merge)
Qst_Mode_Merge_filt$condition = factor(paste(Qst_Mode_Merge_filt$MatOrPat, Qst_Mode_Merge_filt$mode, sep = '_'))

# For the maternal and paternal only conditions (Figure 5C)
p10 <- ggplot(filter(Qst_Mode_Merge_filt, generation == 4989), mapping=aes(Qst,SexRatio,color=condition, fill = 'white')) +
  geom_abline(slope = -0.5, intercept = 0.5, linewidth =1) +
  geom_errorbarh(aes(xmin=Qst-ci.x, xmax=Qst+ci.x), height=.005, size=1) +
  geom_errorbar(aes(ymin=SexRatio-ci.y, ymax=SexRatio+ci.y), width=.03, size=1) +
  stat_summary(fun.y=mean, geom="point", stroke=2, size=3.5, fill='white', aes(shape = condition, color=condition)) +
  xlim(0,1) +
  ylim(0,0.5) +
  scale_shape_manual( values = c(21, 21)) +
  scale_color_manual(values = c('#D9911A', '#3305A3')) +
  ylab("Proportion Male") +
  xlab("Qst") +
  theme_classic() +
  theme(axis.text=element_text(size=20),
        axis.title=element_text(size=25,face="bold"), strip.text = element_text(size = 18),
        legend.position = "none", panel.border = element_rect(colour = "black", fill=NA, linewidth=2), panel.grid.major = element_line(color = 'lightgrey', linetype = 'dotted'))

p10


#For simulations with both maternal and paternal loci (Figure 5E)

Qst_Mode_Merge_filt = rbind(c2_Merge,c4_Merge)
Qst_Mode_Merge_filt$condition = factor(paste(Qst_Mode_Merge_filt$MatOrPat, Qst_Mode_Merge_filt$mode, sep = '_'))

p11 <- ggplot(filter(Qst_Mode_Merge_filt, generation == 4989), mapping=aes(Qst,SexRatio,color=condition, fill = 'white')) +
  geom_abline(slope = -0.5, intercept = 0.5, linewidth =1) +
  geom_errorbarh(aes(xmin=Qst-ci.x, xmax=Qst+ci.x), height=.005, size=1) +
  geom_errorbar(aes(ymin=SexRatio-ci.y, ymax=SexRatio+ci.y), width=.03, size=1) +
  stat_summary(fun.y=mean, geom="point", stroke=2, size=3.5, fill='white', aes(shape = condition, color=condition)) +
  xlim(0,1) +
  ylim(0,0.5) +
  scale_shape_manual( values = c(21, 21)) +
  scale_color_manual(values = c('#D9911A', '#3305A3')) +
  ylab("Proportion Male") +
  xlab("Qst") +
  theme_classic() +
  theme(axis.text=element_text(size=20),
        axis.title=element_text(size=25,face="bold"), strip.text = element_text(size = 18),
        legend.position = "none", panel.border = element_rect(colour = "black", fill=NA, linewidth=2), panel.grid.major = element_line(color = 'lightgrey', linetype = 'dotted'))

p11


#This section is for discrete generations

SR_Disc %>% pivot_longer(cols=c('QstPat', 'QstMat'),
                         names_to='MatOrPat',
                         values_to='Qst') -> Qst_Disc

Qst_Disc_SE <- summarySE(Qst_Disc, measurevar="Qst", groupvars=c("generation","nFounders_char", "mode", "MatOrPat"))

c1 = Qst_Disc_SE[Qst_Disc_SE$mode == 'F' &  Qst_Disc_SE$MatOrPat == 'QstMat',]
c2 = Qst_Disc_SE[Qst_Disc_SE$mode == 'F&M' &  Qst_Disc_SE$MatOrPat == 'QstMat',]
c3 = Qst_Disc_SE[Qst_Disc_SE$mode == 'M' &  Qst_Disc_SE$MatOrPat == 'QstPat',]
c4 = Qst_Disc_SE[Qst_Disc_SE$mode == 'F&M' &  Qst_Disc_SE$MatOrPat == 'QstPat',]

Qst_Disc_SE_filt = rbind(c1,c3)
Qst_Disc_SE_filt$condition = factor(paste(Qst_Disc_SE_filt$MatOrPat, Qst_Disc_SE_filt$mode, sep = '_'))
Qst_Disc_SE_filt = na.omit(Qst_Disc_SE_filt)
Qst_Disc_SE_filt$condition = droplevels(Qst_Disc_SE_filt$condition)

#plot Qst by maternal and paternal effects in the maternal and paternal only conditions
p12 <- ggplot(Qst_Disc_SE_filt, aes(x=generation, y=Qst, color=condition, shape=condition,fill='white')) + 
  geom_errorbar(aes(ymin=Qst-ci, ymax=Qst+ci),width=0,size=1) +
  geom_point(aes(stroke=1),size=1) +
  scale_x_continuous( limits=c(0,5000)) +
  theme_classic() +
  ylab("Qst") +
  xlab("Generation") +
  ylim(0,1) +
  theme(axis.text=element_text(size=20),
        axis.title=element_text(size=25,face="bold"), strip.background =element_rect(fill="white"), strip.text = element_text(size = 25),
        legend.position = "none", panel.border = element_rect(colour = "black", fill=NA, linewidth=2), panel.grid.major = element_line(color = 'lightgrey', linetype = 'dotted')) + 
  scale_shape_manual(values = c(21, 21)) +
  scale_color_manual(values = c('#D9911A', '#3305A3')) +
  scale_fill_manual(values ='white')

wrapped_p12 <- p12 + facet_wrap(~nFounders_char)
wrapped_p12


#plot Qst by maternal and paternal effects with 5 founders for (Figure 5B)
p13 <- ggplot(filter(Qst_Disc_SE_filt, nFounders_char == "5 Founders"), aes(x=generation, y=Qst, color=condition, shape=condition,fill='white')) + 
  geom_errorbar(aes(ymin=Qst-ci, ymax=Qst+ci),width=0,size=1) +
  geom_point(aes(stroke=1),size=1) +
  scale_x_continuous( limits=c(0,5000)) +
  theme_classic() +
  ylab("Qst") +
  xlab("Generation") +
  ylim(0,1) +
  theme(axis.text=element_text(size=20),
        axis.title=element_text(size=25,face="bold"), strip.background =element_rect(fill="white"), strip.text = element_text(size = 25),
        legend.position = "none", panel.border = element_rect(colour = "black", fill=NA, linewidth=2), panel.grid.major = element_line(color = 'lightgrey', linetype = 'dotted')) + 
  scale_shape_manual(values = c(21, 21)) +
  scale_color_manual(values = c('#D9911A', '#3305A3')) +
  scale_fill_manual(values ='white')

wrapped_p13 <- p13 + facet_wrap(~nFounders_char)
wrapped_p13

# plot sex ratio by Qst and by founder in the final generation

Qst_Disc_SR_SE <- summarySE(Qst_Disc, measurevar="SexRatio", groupvars=c("generation","nFounders_char", "mode", "MatOrPat"))
Qst_Disc_Merge <- merge(Qst_Disc_SE,Qst_Disc_SR_SE,by=c("generation","nFounders_char","mode","MatOrPat"))

c1_Merge  = Qst_Disc_Merge[Qst_Disc_Merge$mode == 'F' &  Qst_Disc_Merge$MatOrPat == 'QstMat',]
c2_Merge  = Qst_Disc_Merge[Qst_Disc_Merge$mode == 'F&M' &  Qst_Disc_Merge$MatOrPat == 'QstMat',]
c3_Merge  = Qst_Disc_Merge[Qst_Disc_Merge$mode == 'M' &  Qst_Disc_Merge$MatOrPat == 'QstPat',]
c4_Merge  = Qst_Disc_Merge[Qst_Disc_Merge$mode == 'F&M' &  Qst_Disc_Merge$MatOrPat == 'QstPat',]

Qst_Disc_Merge_filt = rbind(c1_Merge,c3_Merge)
Qst_Disc_Merge_filt$condition = factor(paste(Qst_Disc_Merge_filt$MatOrPat, Qst_Disc_Merge_filt$mode, sep = '_'))

# For the maternal and paternal only conditions (Figure 5C)
p14 <- ggplot(filter(Qst_Disc_Merge_filt, generation == 4989), mapping=aes(Qst,SexRatio,color=condition, fill = 'white')) +
  geom_abline(slope = -0.5, intercept = 0.5, linewidth =1) +
  geom_errorbarh(aes(xmin=Qst-ci.x, xmax=Qst+ci.x), height=.005, size=1) +
  geom_errorbar(aes(ymin=SexRatio-ci.y, ymax=SexRatio+ci.y), width=.03, size=1) +
  stat_summary(fun.y=mean, geom="point", stroke=2, size=3.5, fill='white', aes(shape = condition, color=condition)) +
  xlim(0,1) +
  ylim(0,0.5) +
  scale_shape_manual( values = c(21, 21)) +
  scale_color_manual(values = c('#D9911A', '#3305A3')) +
  ylab("Proportion Male") +
  xlab("Qst") +
  theme_classic() +
  theme(axis.text=element_text(size=20),
        axis.title=element_text(size=25,face="bold"), strip.text = element_text(size = 18),
        legend.position = "none", panel.border = element_rect(colour = "black", fill=NA, linewidth=2), panel.grid.major = element_line(color = 'lightgrey', linetype = 'dotted'))

p14

#For simulations with both maternal and paternal loci (Figure 5E)

Qst_Disc_Merge_filt = rbind(c2_Merge,c4_Merge)
Qst_Disc_Merge_filt$condition = factor(paste(Qst_Disc_Merge_filt$MatOrPat, Qst_Disc_Merge_filt$mode, sep = '_'))

p14 <- ggplot(filter(Qst_Disc_Merge_filt, generation == 4989), mapping=aes(Qst,SexRatio,color=condition, fill = 'white')) +
  geom_abline(slope = -0.5, intercept = 0.5, linewidth =1) +
  geom_errorbarh(aes(xmin=Qst-ci.x, xmax=Qst+ci.x), height=.005, size=1) +
  geom_errorbar(aes(ymin=SexRatio-ci.y, ymax=SexRatio+ci.y), width=.03, size=1) +
  stat_summary(fun.y=mean, geom="point", stroke=2, size=3.5, fill='white', aes(shape = condition, color=condition)) +
  xlim(0,1) +
  ylim(0,0.5) +
  scale_shape_manual( values = c(21, 21)) +
  scale_color_manual(values = c('#D9911A', '#3305A3')) +
  ylab("Proportion Male") +
  xlab("Qst") +
  theme_classic() +
  theme(axis.text=element_text(size=20),
        axis.title=element_text(size=25,face="bold"), strip.text = element_text(size = 18),
        legend.position = "none", panel.border = element_rect(colour = "black", fill=NA, linewidth=2), panel.grid.major = element_line(color = 'lightgrey', linetype = 'dotted'))

p14

#How much of the variance in sex ratio is captured by Qst for overlapping generations? 

Qst_Mode$condition = factor(paste(Qst_Mode$MatOrPat, Qst_Mode$mode, sep = '_'))


Qst_Mode_lm=lm(data = filter(Qst_Mode, generation == 4989, condition %in% c("QstMat_F", "QstPat_M")), SexRatio ~  Qst)


summary(Qst_Mode_lm)

"""
Residuals:
  Min        1Q    Median        3Q       Max 
-0.060822 -0.014710  0.000326  0.014295  0.065270 

Coefficients:
  Estimate Std. Error t value Pr(>|t|)    
(Intercept)  0.490470   0.002520  194.61   <2e-16 ***
  Qst         -0.272022   0.004601  -59.12   <2e-16 ***
  ---
  Signif. codes:  0 ‘***’ 0.001 ‘**’ 0.01 ‘*’ 0.05 ‘.’ 0.1 ‘ ’ 1

Residual standard error: 0.02194 on 448 degrees of freedom
Multiple R-squared:  0.8864,	Adjusted R-squared:  0.8861 
F-statistic:  3495 on 1 and 448 DF,  p-value: < 2.2e-16

"""

#How much of the variance in sex ratio is captured by Qst for discrete generations? 

Qst_Disc$condition = factor(paste(Qst_Disc$MatOrPat, Qst_Disc$mode, sep = '_'))

Qst_Disc_lm=lm(data = filter(Qst_Disc, generation == 4989, condition %in% c("QstMat_F", "QstPat_M")), SexRatio ~  Qst)

summary(Qst_Disc_lm)


"""
Residuals:
      Min        1Q    Median        3Q       Max 
-0.064286 -0.012638  0.000081  0.013099  0.077614 

Coefficients:
             Estimate Std. Error t value Pr(>|t|)    
(Intercept)  0.456848   0.002368  192.88   <2e-16 ***
Qst         -0.227517   0.004380  -51.95   <2e-16 ***
---
Signif. codes:  0 ‘***’ 0.001 ‘**’ 0.01 ‘*’ 0.05 ‘.’ 0.1 ‘ ’ 1

Residual standard error: 0.02104 on 448 degrees of freedom
Multiple R-squared:  0.8576,	Adjusted R-squared:  0.8573 
F-statistic:  2698 on 1 and 448 DF,  p-value: < 2.2e-16
"""

########################################
### Mating After Dispersal in a Pool ###
########################################

#plot sex ratio in final generation (Figure 6, middle)
p15 <- ggplot(filter(SR_AD, generation == 2591 | generation == 2590 | generation == 2589 | generation == 2588), mapping=aes(factor(nFounders),SexRatio,fill=mode)) + 
  geom_boxplot(alpha=0.85,position = "dodge", outlier.shape = NA) +
  ylim(.15,0.56) +
  scale_fill_manual(values = c('#D9911A', '#666666','#3305A3')) +
  ylab("Proportion Male") +
  xlab("Founder Number") +
  theme_classic() +
  theme(axis.text=element_text(size=20),
        axis.title=element_text(size=25,face="bold"), strip.text = element_text(size = 18),
        legend.position = "none", panel.border = element_rect(colour = "black", fill=NA, linewidth=2), panel.grid.major = element_line(color = 'lightgrey', linetype = 'dotted'))


wrapped_p15 <- p15 + facet_grid(~G,  scales = "free", space='free')

wrapped_p15


#plot genetic effects in the both condition

SR_AD %>% pivot_longer(cols=c('matGE', 'patGE'),
                              names_to='MatOrPat',
                              values_to='GE') -> GE_AD


#plot genetic effects in final generation (Supplemental figure 1, middle)
p16 <- ggplot(filter(GE_AD, generation == 2591 | generation == 2590 | generation == 2589 | generation == 2588, mode == 3), mapping=aes(factor(nFounders),GE,fill=MatOrPat)) + 
  geom_boxplot(alpha=0.85,position = "dodge", outlier.shape = NA) +
  ylim(-2,3) +
  scale_fill_manual(values = c('#D9911A','#3305A3')) +
  ylab("Genetic Effect") +
  xlab("Founder Number") +
  theme_classic() +
  theme(axis.text=element_text(size=20),
        axis.title=element_text(size=25,face="bold"), strip.text = element_text(size = 18),
        legend.position = "none", panel.border = element_rect(colour = "black", fill=NA, linewidth=2), panel.grid.major = element_line(color = 'lightgrey', linetype = 'dotted'))


wrapped_p16 <- p16 + facet_grid(~G,  scales = "free", space='free')

wrapped_p16

# Is there a difference in the evolved sex ratios between modes for one-generation patches?

lm1_AD_G1=lm(data = filter(SR_AD,  generation == 2591), formula = SexRatio ~ mode + nFounders)
lm2_AD_G1=lm(data = filter(SR_AD,  generation == 2591), formula = SexRatio ~ nFounders)

anova(lm1_AD_G1, lm2_AD_G1, test = "Chisq")

"""
Analysis of Variance Table

Model 1: SexRatio ~ mode + as.factor(nFounders)
Model 2: SexRatio ~ as.factor(nFounders)
  Res.Df     RSS Df   Sum of Sq Pr(>Chi)
1    293 0.11043                        
2    295 0.11060 -2 -0.00016597   0.8024
"""

# Is there a difference for evolved sex ratios between modes for founder number = 1 across all generations in a patch?

lm1_AD_LMC=lm(data = filter(SR_AD,  generation == 2591 | generation == 2590 | generation == 2589 | generation == 2588, nFounders == 1), formula = SexRatio ~ mode + G)
lm2_AD_LMC=lm(data = filter(SR_AD,  generation == 2591 | generation == 2590 | generation == 2589 | generation == 2588, nFounders == 1), formula = SexRatio ~  G)

anova(lm1_AD_LMC, lm2_AD_LMC, test = "Chisq")

"""
Analysis of Variance Table

Model 1: SexRatio ~ mode + G
Model 2: SexRatio ~ G
  Res.Df      RSS Df  Sum of Sq Pr(>Chi)  
1    234 0.099576                         
2    236 0.102458 -2 -0.0028817  0.03385 *
"""

# Is there a difference for evolved sex ratios between modes across all generations in a patch?

lm1_AD_byG=lm(data = filter(SR_AD,  generation == 2591 | generation == 2590 | generation == 2589 | generation == 2588), formula = SexRatio ~ mode + G + nFounders)
lm2_AD_byG=lm(data = filter(SR_AD,  generation == 2591 | generation == 2590 | generation == 2589 | generation == 2588), formula = SexRatio ~ G + nFounders)

anova(lm1_AD_byG, lm2_AD_byG, test = "Chisq")

"""
Analysis of Variance Table

Model 1: SexRatio ~ mode + G + nFounders
Model 2: SexRatio ~ G + nFounders
  Res.Df     RSS Df Sum of Sq Pr(>Chi)    
1   1190 0.70408                          
2   1192 0.73569 -2 -0.031608 2.51e-12 ***
---
Signif. codes:  0 ‘***’ 0.001 ‘**’ 0.01 ‘*’ 0.05 ‘.’ 0.1 ‘ ’ 1
"""

###############################
### Mating Before Dispersal ###
###############################


#plot sex ratio in final generation (Figure 6, bottom)
p17 <- ggplot(filter(SR_BD,  generation == 2592 | generation == 2591 | generation == 2590 | generation == 2589), mapping=aes(factor(nFounders),SexRatio,fill=mode)) + 
  geom_boxplot(alpha=0.85,position = "dodge", outlier.shape = NA) +
  ylim(.15,0.56) +
  scale_fill_manual(values = c('#D9911A', '#666666','#3305A3')) +
  ylab("Proportion Male") +
  xlab("Founder Number") +
  theme_classic() +
  theme(axis.text=element_text(size=20),
        axis.title=element_text(size=25,face="bold"), strip.text = element_text(size = 18),
        legend.position = "none", panel.border = element_rect(colour = "black", fill=NA, linewidth=2), panel.grid.major = element_line(color = 'lightgrey', linetype = 'dotted'))


wrapped_p17 <- p17 + facet_grid(~G,  scales = "free", space='free')

wrapped_p17


#plot genetic effects in the both condition

SR_BD %>% pivot_longer(cols=c('matGE', 'patGE'),
                              names_to='MatOrPat',
                              values_to='GE') -> GE_BD


#plot genetic effects in final generation (Supplemental figure 1, bottom)
p18 <- ggplot(filter(GE_BD, generation == 2592 | generation == 2591 | generation == 2590 | generation == 2589, mode == 3), mapping=aes(factor(nFounders),GE,fill=MatOrPat)) + 
  geom_boxplot(alpha=0.85,position = "dodge", outlier.shape = NA) +
  ylim(-2,3) +
  scale_fill_manual(values = c('#D9911A','#3305A3')) +
  ylab("Genetic Effect") +
  xlab("Founder Number") +
  theme_classic() +
  theme(axis.text=element_text(size=20),
        axis.title=element_text(size=25,face="bold"), strip.text = element_text(size = 18),
        legend.position = "none", panel.border = element_rect(colour = "black", fill=NA, linewidth=2), panel.grid.major = element_line(color = 'lightgrey', linetype = 'dotted'))


wrapped_p18 <- p18 + facet_grid(~G,  scales = "free", space='free')

wrapped_p18


# Is there a difference in the evolved sex ratios between modes for one-generation patches?

lm1_BD_G1=lm(data = filter(SR_BD,  generation == 2592), formula = SexRatio ~ mode + nFounders)
lm2_BD_G1=lm(data = filter(SR_BD,  generation == 2592), formula = SexRatio ~  nFounders)

anova(lm1_BD_G1, lm2_BD_G1, test = "Chisq")


"""
Analysis of Variance Table

Model 1: SexRatio ~ as.factor(mode) + as.factor(nFounders)
Model 2: SexRatio ~ as.factor(nFounders)
  Res.Df      RSS Df   Sum of Sq Pr(>Chi)
1    293 0.098192                        
2    295 0.098264 -2 -7.1547e-05   0.8988
"""

# Is there a difference for evolved sex ratios for founder number = 1 across all generations in a patch?


lm1_BD_F1=lm(data = filter(SR_BD,  generation == 2592 | generation == 2591 | generation == 2590 | generation == 2589, nFounders == 1), formula = SexRatio ~ mode + G)
lm2_BD_F1=lm(data = filter(SR_BD,  generation == 2592 | generation == 2591 | generation == 2590 | generation == 2589, nFounders == 1), formula = SexRatio ~  G)

anova(lm1_BD_F1, lm2_BD_F1, test = "Chisq")


"""
Analysis of Variance Table

Model 1: SexRatio ~ mode + G
Model 2: SexRatio ~ G
  Res.Df      RSS Df   Sum of Sq Pr(>Chi)
1    234 0.085417                        
2    236 0.085505 -2 -8.7977e-05   0.8865
"""

# Is there a difference for evolved sex ratios across all generations in a patch?

lm1_BD_byG=lm(data = filter(SR_BD,  generation == 2592 | generation == 2591 | generation == 2590 | generation == 2589), formula = SexRatio ~ mode + G + nFounders)
lm2_BD_byG=lm(data = filter(SR_BD,  generation == 2592 | generation == 2591 | generation == 2590 | generation == 2589), formula = SexRatio ~ G + nFounders)

anova(lm1_BD_byG, lm2_BD_byG, test = "Chisq")

"""
Analysis of Variance Table

Model 1: SexRatio ~ mode + G + nFounders
Model 2: SexRatio ~ G + nFounders
  Res.Df     RSS Df Sum of Sq  Pr(>Chi)    
1   1190 0.60484                           
2   1192 0.69580 -2 -0.090958 < 2.2e-16 ***
---
Signif. codes:  0 ‘***’ 0.001 ‘**’ 0.01 ‘*’ 0.05 ‘.’ 0.1 ‘ ’ 1
"""

###############################################
### Mating After Dispersal and Colonization ###
###############################################


#plot sex ratio in final generation (Figure 6, top)
p19 <- ggplot(filter(SR_MAC,  generation == 2591 | generation == 2590 | generation == 2589 | generation == 2588), mapping=aes(factor(nFounders),SexRatio,fill=mode)) + 
  geom_boxplot(alpha=0.85,position = "dodge", outlier.shape = NA) +
  ylim(.15,0.56) +
  scale_x_discrete(limits=factor(c(1,2,3,4,5)), drop=FALSE) +
  scale_fill_manual(values = c('#D9911A', '#666666','#3305A3')) +
  ylab("Proportion Male") +
  xlab("Founder Number") +
  theme_classic() +
  theme(axis.text=element_text(size=20),
        axis.title=element_text(size=25,face="bold"), strip.text = element_text(size = 18),
        legend.position = "none", panel.border = element_rect(colour = "black", fill=NA, linewidth=2), panel.grid.major = element_line(color = 'lightgrey', linetype = 'dotted'))


wrapped_p19 <- p19 + facet_grid(~G,  scales = "free", space='free')

wrapped_p19

#plot genetic effects in the both condition 

SR_MAC %>% pivot_longer(cols=c('matGE', 'patGE'),
                              names_to='MatOrPat',
                              values_to='GE') -> GE_MAC

#plot genetic effects in final generation (Supplemental figure 1, top)
p20 <- ggplot(filter(GE_MAC, generation == 2591 | generation == 2590 | generation == 2589 | generation == 2588, mode == 3), mapping=aes(factor(nFounders),GE,fill=MatOrPat)) + 
  geom_boxplot(alpha=0.85,position = "dodge", outlier.shape = NA) +
  ylim(-2,3) +
  scale_x_discrete(limits=factor(c(1,2,3,4,5)), drop=FALSE) +
  scale_fill_manual(values = c('#D9911A','#3305A3')) +
  ylab("Genetic Effect") +
  xlab("Founder Number") +
  theme_classic() +
  theme(axis.text=element_text(size=20),
        axis.title=element_text(size=25,face="bold"), strip.text = element_text(size = 18),
        legend.position = "none", panel.border = element_rect(colour = "black", fill=NA, linewidth=2), panel.grid.major = element_line(color = 'lightgrey', linetype = 'dotted'))


wrapped_p20 <- p20 + facet_grid(~G,  scales = "free", space='free')

wrapped_p20


# Is there a difference in the evolved sex ratios for one generation patches?

lm1_MAC_G1=lm(data = filter(SR_MAC,  generation == 2591), formula = SexRatio ~ mode + nFounders)
lm2_MAC_G1=lm(data = filter(SR_MAC,  generation == 2591), formula = SexRatio ~  nFounders)

anova(lm1_MAC_G1, lm2_MAC_G1, test = "Chisq")

"""
Analysis of Variance Table

Model 1: SexRatio ~ mode + nFounders
Model 2: SexRatio ~ nFounders
  Res.Df     RSS Df  Sum of Sq Pr(>Chi)
1    175 0.14650                       
2    177 0.14829 -2 -0.0017904   0.3432
"""

# Is there a difference for evolved sex ratios for founder number = 1 across all generations in a patch?

lm1_MAC_F1=lm(data = filter(SR_MAC,  generation == 2591 | generation == 2590 | generation == 2589 | generation == 2588, nFounders == 2), formula = SexRatio ~ mode + G)
lm2_MAC_F1=lm(data = filter(SR_MAC,  generation == 2591 | generation == 2590 | generation == 2589 | generation == 2588, nFounders == 2), formula = SexRatio ~  G)

anova(lm1_MAC_F1, lm2_MAC_F1, test = "Chisq")

"""
Analysis of Variance Table

Model 1: SexRatio ~ mode + G
Model 2: SexRatio ~ G
  Res.Df     RSS Df   Sum of Sq Pr(>Chi)
1    175 0.15576                        
2    177 0.15621 -2 -0.00044177   0.7802
"""

# Is there a difference for evolved sex ratios across all generations in a patch?

lm1_MAC_byG=lm(data = filter(MAC_N20,  generation == 2591 | generation == 2590 | generation == 2589 | generation == 2588), formula = SexRatio ~ mode + G + nFounders)
lm2_MAC_byG=lm(data = filter(MAC_N20,  generation == 2591 | generation == 2590 | generation == 2589 | generation == 2588), formula = SexRatio ~ G + nFounders)

anova(lm1_MAC_byG, lm2_MAC_byG, test = "Chisq")

"""
Analysis of Variance Table

Model 1: SexRatio ~ mode + G + nFounders
Model 2: SexRatio ~ G + nFounders
  Res.Df    RSS Df Sum of Sq  Pr(>Chi)    
1    896 1.5412                           
2    897 1.5733 -1 -0.032053 1.583e-05 ***
"""
```
