## Supplementary material for "Sexual antagonism evolves when autosomes influence offspring sex ratio": SlimWrapper.py

```
from argparse import ArgumentParser
import subprocess
import os, sys

def parse_rel(slim_string):
    """
    Parse the output of SLiM to extract whatever data we're looking for.
    If we want to do a more complex analysis on the output of the SLiM file,
    this is where we do it.
    Args:
        slim_string: the entire output of a run of SLiM.
    Return
        output: the desired output we want from the SLiM simulation.
    """
    # The example SLiM file has been configured such that all the
    # output we want is printed on lines that start with "OUT:"
    # so we'll discard all other output lines."
    output = ""
    lines = slim_string.split('\n')
    for line in lines:
        if line.startswith("OUT:"):
            output += '\n' + line.split(":")[1]  # Discard the "OUT:" from the output line.
    return output

def run_slim(command_line_args):
    """
    Runs SLiM using subprocess.
    Args:
        command_line_args: list; a list of command line arguments.
    return: The entire SLiM output as a string.
    """
    slim = subprocess.Popen(command_line_args,
                            stdout=subprocess.PIPE,
                            stderr=subprocess.PIPE,
                            universal_newlines=True)
    out, err = slim.communicate()
    # For debugging purposes:
    # std.out from the subprocess is in slim.communicate()[0]
    # std.error from the subprocess is in slim.communicate()[1]
    # Errors from the process can be printed with:
    print(err)
    return out


def configure_slim_command_line(args_dict):
    """
    Sets up a list of command line arguments for running SLiM.
    Args:
        args_dict: a dictionary of arg parser arguments.
    Return
        clargs: A formated list of the arguments.
    """
    # We're running SLiM, so the first arg is simple:
    clargs = "slim "
    # The filename of the source file must be the last argument:
    source = args_dict.pop("source")
    # Add each argument from arg parser to the command line arguemnts for SLiM

    for arg in args_dict:
        if isinstance(args_dict[arg], bool):
            clargs += f"-d {arg}={'T' if args_dict[arg] else 'F'} "
        else:
            clargs += f"-d {arg}={args_dict[arg]} "
    # Add the source file, and return the string split into a list.
    clargs += source
    return clargs.split()

def main():
    """
    print(slim_result)


    1. Configure using argparse.
    2. Generate the command line list to pass to subprocess through the run_slim() function.
    3. Run SLiM.
    4. Process the output of SLiM to extract the information we want.
    5. Print the results.
    """

    current_wd = os.getcwd()


    # Get args from arg parser:
    parser = ArgumentParser()
    parser.add_argument('-src', '--source', default="males_as_agents.slim", type=str,
                        help=r"SLiM file to be run. Default 'males_as_agents.slim'")
    parser.add_argument('-header', '--print_header', action='store_true', default=False,
                        help='If this is set, python prints a header for a csv file.')

    # The all caps names of the following arguments must exactly match
    # the names of the constants we want to define in SLiM.
    parser.add_argument('-PopSR', '--PopSR', default=0.5, type=float,
                        help='Starting sex ratio as proportion of males produced')
    parser.add_argument('-broodSize', '--broodSize', default=10, type=int,
                        help='number of offspring for a single female per generation')
    parser.add_argument('-N', '--N', default=200, type=int,
                        help='number of subpopulations default is 20')
    parser.add_argument('-m', '--m', default=5, type=int,
                        help='number of founders per migration event, cannot be lower than 2')
    parser.add_argument('-R', '--R', default=3e-6, type=int,
                        help='recombination rate per basepair (averege one per chromosome), default is 6e-6')
    parser.add_argument('-mode', '--mode', default=1, type=int,
            help='Where the sex ratio QTL are active, can be in maternal:1, paternal:2, or both:3')
    parser.add_argument('-G', '--G', default=3, type=int,
            help='Number of generations before migration.')


    args_dict = vars(parser.parse_args())

    # The '-header' argument prints a header for the output. This can
    # help generate a nice CSV by adding this argument to the first SLiM run:
    if args_dict.pop("print_header", None):
        print("nFounders\G\tmode\tPopSR\tgeneration\tQstMat\tQstPat\tSexRatio\tpatGE\tmatGE")

    # Next, assemble the command line arguments in the way we want to for SLiM:
    clargs = configure_slim_command_line(args_dict)

    

    # Run the file with the desired arguments.
    slim_result = run_slim(clargs)

    #Parse and analyze the result.
    parsed_rel = parse_rel(slim_result)

    print(parsed_rel)

if __name__ == "__main__":
    main()
```
